# Dystrophic changes of nigrostriatal axons harboring a Synj1 Parkinson mutation suggest catastrophic failure of endocytic mechanisms

**DOI:** 10.64898/2026.06.24.733515

**Authors:** Yumei Wu, Peng Xu, James Moran, C. Shan Xu, Kenneth J. Hayworth, Mian Cao, Lin Shao, D. James Surmeier, Harald F. Hess, Pietro De Camilli

## Abstract

Synaptojanin 1 is a brain enriched phosphoinositide phosphatase implicated in endocytosis at the synapse. A mutation (R258Q) that selectively impairs its Sac1 phosphatase domain causes early onset familial Parkinsonism. Neurons of mice with this mutation display synaptic vesicle traffic defects across the brain, but selective dystrophic changes in a subset of dopaminergic axons in the dorsolateral striatum. Using correlative light microscopy-FIB-SEM of mutant mouse striata to visualize in 3D these abnormal structures we show that they represent clusters of focal axonal dilations harboring massive, onion-like DAT enriched plasma membrane infoldings, generally localized next to cell bodies of neighboring cells, often engulfing evaginations of such cells. This dysmorphia was associated with a deficit in dopamine release in the same striatal region. Given the involvement of Synj1 in endocytic mechanisms, these structures may reflect an imbalance between exocytosis and endocytosis. Their occurrence only in a subset of axons suggest a vulnerability threshold of these axons beyond which the expansion of the plasma membrane is not counteracted by compensatory mechanisms.

---

Human genetic studies have identified more than 20 genes whose mutations are responsible for Parkinson’s disease (1–3). A key priority is to build on this information to elucidate disease mechanisms. One of these genes, *PARK20*, encodes Synaptojanin 1 (Synj1), a PI(4,5)P_2_ phosphatase enriched in presynaptic nerve terminals where it achieves its function through two distinct, tandemly arranged phosphatase domains - a Sac1 domain (primarily a 4-phosphatase domain) and a 5-phosphatase domain - which cleave sequentially the phosphates at the 5 and 4 positions of the inositol ring of this phospholipid (4–6). One of the major and best known roles of Synj1 at synapses is to mediate the shedding of clathrin coats and other endocytic factors during synaptic vesicle endocytosis (5, 7–9), although additional roles have been reported (6, 10–12). Mutations resulting in absence or nearly complete absence of Synj1 result in early postnatal lethality in mice (5) and severe childhood disorders in humans (13, 14).

In 2013, two independent studies reported that a homozygous mutation that selectively impairs the catalytic activity of its Sac1 domain (R258Q) causes recessive familial early-onset Parkinsonism (15, 16). Since then, other *SYNJ1* mutations that only partially impair SYNJ1’s function were found to be associated with PD (17, 18), but the R258Q mutation remains the best characterized. We have shown that neurons of mice homozygous for the corresponding amino acid substitution (R259Q), referred to henceforth as Synj1^RQ^ mice, display neurological manifestations reminiscent of those of human patients. This phenotype is accompanied by endocytic defects at synapses which are not restricted to dopaminergic (DAergic) neurons, consistent with the housekeeping role of Synj1 (12, 19–21). These changes were more prominent at inhibitory synapses, which typically have a high rate of spontaneous activity and thus of synaptic vesicle recycling (19). In addition, dystrophic changes were observed selectively in the dorsal striatum of these mice. They were represented by an accumulation of large abnormal foci of immunoreactivity for markers of DAergic axons, such as the plasma membrane DA transporter (DAT) and tyrosine hydroxylase (TH), which stood out among the normal fine dense network of these axons (19–21). The presence of these foci roughly correlated in abundance with the occurrence in the same brain regions of whorls of multilamellar structures detectable by electron microscopy (EM) (19, 20). However, a correspondence between these two structures has not been proven until now.

The goal of this study was to fill this gap and reveal the precise ultrastructure of these dystrophic axonal changes in order to advance understanding of disease mechanisms. To this end, we generated a Synj1^RQ^ mouse model in which DA neurons express tdTomato, thus making possible correlative light-electron microscopy (CLEM) analysis. We prove that dystrophic changes observed by light microscopy in the dorsal striatum correspond to clusters of onion-like invaginations of the axonal plasma membrane of DA axons which often engulf portions of adjacent cells. In addition, DA release in the same striatal region, particularly with repetitive stimulation, was significantly impaired. In view of the importance of Synj1 in the endocytic pathway, onion-like invaginations may result from a chronic imbalance between addition of membrane to the plasma membrane by exocytosis and removal by endocytosis. As these changes occur only in a subset of axons and axon segments, they indicate that DA axons with impaired Synj1 function may be in a vulnerable state, which in a subset of them can progress to an overt pathological state.

## RESULTS

### Generation of Synj1^RQ^ mice expressing tdTomato in DAergic neurons

To generate mice where the entire DAergic axonal network in the striatum could be selectively visualized by fluorescence microscopy without manipulations that would perturb ultrastructure (such as detergent-based membrane permeabilization required for immunofluorescence), we interbred previously described knock-in mice harboring the Synj1^RQ^ mutation (19), or WT mice as controls, with Ai9 mice which express tdTomato after Cre-dependent recombination (22) and with DAT^IREScre^ mice which express Cre under the control of the plasma membrane dopamine transporter DAT (23). The resulting control and mutant mice (DAT^IRES^cre; Ai9; WT or *Synj1*^RQ/RQ^ mice) (age ranging from 1.5 to 15 months) were anesthetized and transcardially perfused with an aldehyde-based fixative followed by generation of brain vibratome sections and whole brain CLARITY. As expected, inspection of these sections by fluorescence microscopy revealed a bright labeling of DA neurons (Fig. 1A). tdTomato fluorescence decorated all compartments of these neurons, including cell bodies with their dendrites in the Substantia Nigra and Ventral Tegmental Area (VTA), as well as their axons which terminate with a massive highly branched arbor in the Dorsal Striatum, Nucleus Accumbens and Olfactory Tubercle.

**Fig. 1.**
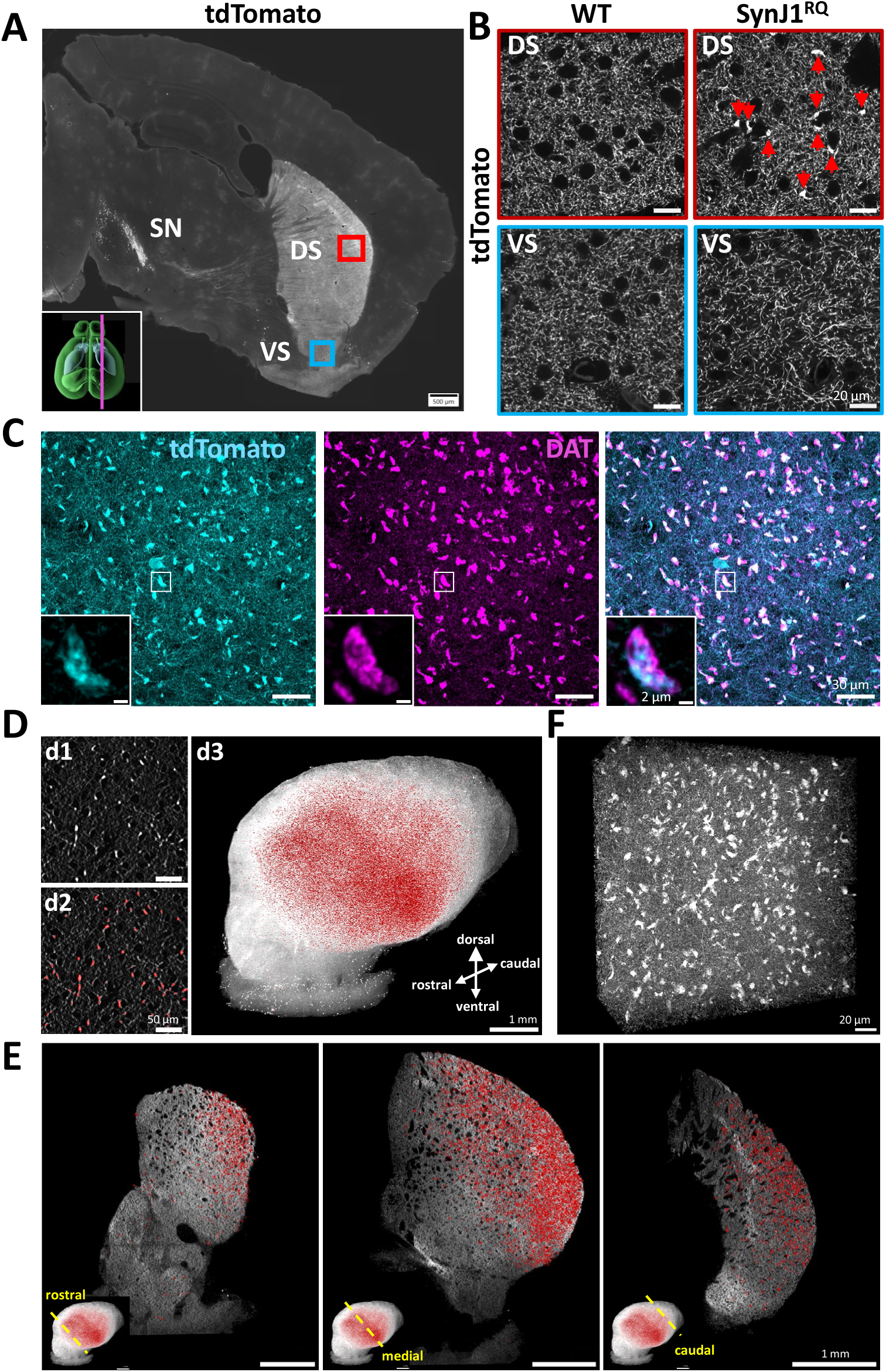
Expression of tdTomato in DAergic neurons reveal presence of dystrophic changes in a subset of their nerve terminals in the dorsal striatum of Synj1^RQ^ mice. **A**. Fluorescence image of a sagittal section of a Synj1^RQ^ mouse (4 month-old, male) expressing tdTomato under the control of a DAT promoter. tdTomato fluorescence (white) labels the entire DAergic neuronal population. SN (Substantia Nigra), DS and VS (Dorsal and Ventral Striatum). The Ventral Tegmental Area (VTA) is not in the plane of the section. **B**. High-power view of the tdTomato fluorescence in the DS and VS (regions outlined by small squares in field A). In all four fields a punctate pattern of tdTomato fluorescence, reflecting the fine arborization of DAergic axons is visible. In addition, large clusters of fluorescence are visible selectively in the DLS of the Synj1^RQ^ mouse brain. **C**. - power view of a region in the DLS of a Synj1^RQ^ striatum (3 month-old, male) showing colocalization of tdTomato fluorescence and DAT immunoreactivity. The inset shows that, when viewed at high magnification, the tdTomato fluorescence (a cytosolic protein) and immunoreactivity for the membrane protein DAT do not precisely overlap as expected. **D.** light-sheet imaging of tdTomato fluorescence in the striatum of a delipidated td-Synj1^RQ^ whole mouse brain (3 month-old, male). **d1** and **d2**. Representative image of a region of the dorsal striatum (d1) and segmentation (red) of the bright tdTomato puncta visible in d1. **d3**. Low maginification side view of the delipidated striatum (visualized by the tdTomato fluorescence, white) showing that segmented dystrophic DAergic axons (red spots) are highly enriched in the DLS. The 3D view is shown in movie S1. **E**. Coronal sections at three different rostro-caudal positions of the striatum in **D** (d3) showing the spatial heterogeneity in the distribution of red tdTomato puncta representing DAergic dystrophic axons (see movie S2). **F.** Confocal microscopy image showing the very high abundance of large bright tdTomato foci, representing dystrophic axons, in a 217×217×100 µm^3^ volume from a region of the DLS most enriched in such foci (see movie S3). Scale bars: **A**, 500 µm; **B**, 20 µm; **C**, 30 µm, inset, 2 µm; D, d1-2, 50 µm, d3, 1 mm; **E**, 1mm; **F**, 20 µm.

### Abnormal foci of tdTomato fluorescence selectively in the dorsolateral striatum (DLS) of Synj1^RQ^ mutant mice

At high-power view, tdTomato fluorescence in the striatum was accounted for by a fine reticulum of axonal arborization (Fig. 1B). In addition, however, sparse larger tdTomato positive foci stood out among the thinner fine axonal reticular network in the dorsal striatum of Synj1^RQ^ mice but not of WT mice. As shown by anti-DAT immunofluorescence, these foci were positive for DAT immunoreactivity (Fig. 1C), confirming that they represent dilations of DAergic axons, in agreement with previous results (20, 24). Thus, these mice represent a good model system to study dystrophic changes induced by the Synj1^RQ^ mutation.

To obtain a global view of the localization of these abnormal dopaminergic axon segments within the striatum, we delipidated the entire brain (25) and then analyzed it by light-sheet microscopy. Large foci of fluorescence were automatically segmented, pseudocolored in red and then superimposed on the total tdTomato fluorescence of the striatum shown in white (Fig. 1D). Three-dimensional views of the entire striatum, which are shown in Fig. 1D (field d3, a side view) and in movie S1, demonstrate the striking and selective accumulation of the bright foci in its dorsolateral region (DorsoLateral Striatum, DLS) and their near absence in the Ventral Striatum (VS). This localization, with a decreasing density of foci in the lateral to medial direction, is further illustrated by virtual optical coronal planes at three different rostro-caudal positions of the striatum in Fig. 1E and movie S2. The high concentration of fluorescent foci in the most lateral region of the DLS is shown in a high-power view of a small region of a 100 µm-thick vibratome section obtained from this region (Fig. 1F and movie S3).

### The Synj1^RQ^ mutation was associated with a deficit in DA release, specifically within the DLS

To determine whether the occurrence of structural changes of DAergic axons in the DLS was accompanied by functional changes, we used the genetically encoded sensor dLight1.3b to explore potential defects of the mutation in DA secretion. dLight1.3b was expressed using a viral vector in the DLS of Synj1^RQ^ mice, and two-photon laser scanning microscopy (2PLSM) was used to estimate electrically evoked DA release. To control for variation in the expression level of the dLight probe, electrically evoked fluorescence measurements were normalized by maximum florescence change (ΔF_max_) evoked by bath application of a saturating concentration of DA (100 µM) at the end of each recording session. The distribution of ΔF_max_ measurements did not differ between genotypes (Fig. S2A), arguing that dLight expression was not affected genotype. In *ex vivo* brain slices from Synj1^RQ^ mice, there was a pronounced reduction in DA release in response to a single stimulus compared to DA release detected in slices of heterozygous littermate controls (*Synj1*^+/RQ^) (39% reduction, *P* = 0.0001) (Fig. 2A). Comparable reductions in release were observed following a train of five stimuli at 10 Hz (30% reduction, *P* < 0.0001) (Fig. 2C). Although the amplitude of the evoked response was reduced in tissue from Synj1^RQ^ mice, the decay kinetics of single-pulse evoked dLight1.3b transients did not differ significantly between genotypes in the DLS [τ: *Synj1^+/RQ^*(control) = 219.3 ms vs *Synj1^RQ^* = 231.4 ms; P = 0.7859] (Fig. S2C–D), suggesting that DA reupake mechanisms were unaffected. To assess the regional specificity of these observations, dLight1.3b was also expressed in the VS. As in the DLS, the distribution of VS ΔF_max_ measurements were similar across genotypes (Fig. S2B). However, in contrast to the DLS, DA release in the VS did not differ significantly In Synj1^RQ^ mice (Fig. S1A-D).

**Fig. 2.**
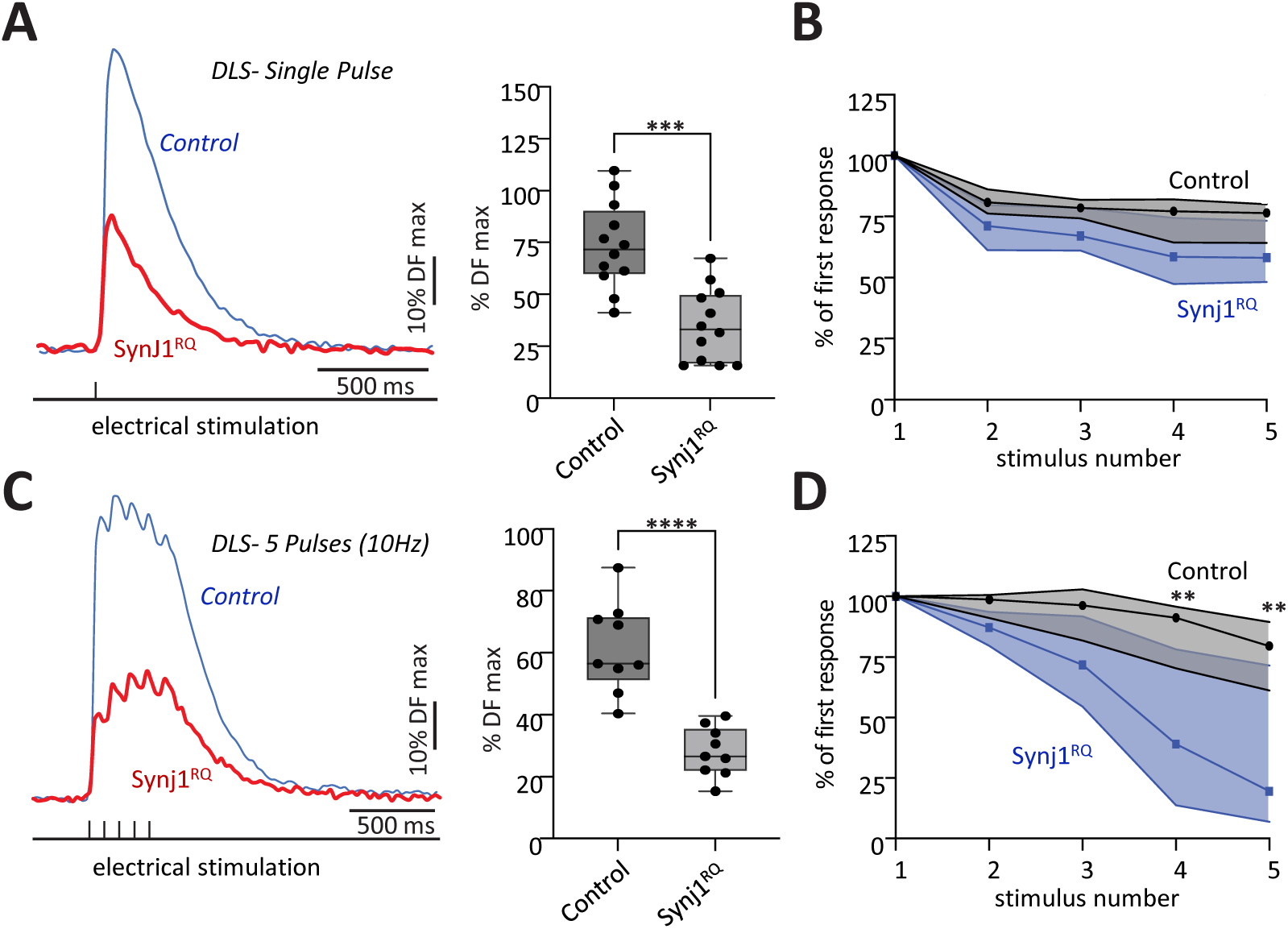
Reduced evoked DA release capacity in the dorsolateral striatum (DLS) of Synj1^RQ^ mice. **A.** Representative traces of DA release as assessed by dLigth1.3b in the DLS evoked from single-pulse stimulation. Traces are normalized to dF max (left). Quantification of the amount of DA released from terminals innervating the DLS in response to single stimulatory events. Quantifications are presented as a percentage of dF max (achieved through bath application of 100 µM DA). Synj1^RQ^: n=12, N=6; *Synj1*^+/RQ^ (control): n=12, N=7. (right). **B.** Plot illustrating mild deficits in response to consecutive single pulses in the DLS of homozygous Synj1^RQ^ mice when release is evoked by a single pulse. Responses are normalized to the first response in the stimulation protocol. Synj1^RQ^: n=11, N=4; *Synj1*^+/RQ^ (control): n=14, N=4.. **C.** Representative trace of DA release in the DLS in response to 5-pulse stimulation at 10 Hz. Traces are normalized to dF max (left). Quantification of the amount of DA released from terminals innervating the DLS in response to a 5-pulse train at 10 Hz. Quantifications are based on the dF of the 5^th^ pulse in the train and are presented as a percentage of dF max. Synj1^RQ^: n=9, N=3; *Synj1*^+/RQ^ (control): n=9, N=4. (right). **D.** Plot illustrating profound deficits in vesicle recycling in the DLS of *Synj1*^RQ^ mice when release is evoked a 5-pulse train at 10 Hz. Responses are normalized to the first response in the stimulation protocol. Synj1^RQ^: n=11, N=3; *Synj1*^+/RQ^ (control): n=11, N=4.

The electrical stimulation (either a single pulse or a five-pulse burst) was also repeated at ten second intervals and DA release was monitored using dLight. In slices from Synj1^RQ^ mice, there was a pronounced, progressive drop in DA release evoked in response to the burst stimulation, in contrast to the response in slices from heterozygous controls (*Synj1*^+/RQ^) (Fig. 2D). The release evoked by a single stimulus trended in the same direction but did not reach statistical significance relative to *Synj1*^+/RQ^ controls (Fig. 2B). No difference was observed in the Nucleus Accumbens (NAc) of the VS in response to the same stimulation patterns (Fig. S2B and D). However, the decay kinetics of evoked DA transients were slower in the VS of Synj1^RQ^ animals relative to heterozygous controls [τ: *Synj1^+/RQ^* (control) = 223.8 ms vs. Synj1^RQ^ = 290.6 ms; P < 0.0127](Fig. S2E–F), indicating that while release amplitude is preserved, DA clearance was impaired in this region. The reason for the discrepancy in this parameter between DLS and VS deserves further investigation. Taken together, these experiments demonstrate that the Synj1 mutation results in a DLS-specific deficit in DA release – paralleling the region-specific abnormality in axonal morphology.

### Dystrophic axonal structures are generally adjacent to cell bodies

Dystrophic tdTomato- and DAT-positive axonal foci in the DLS were generally adjacent to cell bodies and blood vessels (Fig. 3A and 3B). These included cell bodies of spiny projection neurons (SPNs) (DARPP32 immunoreactive) (26), of astrocytes (S100b immunoreactive) (27, 28) and of microglia (Iba1 immunoreactive) (29), thus ruling out specificity for a given cell type (Fig. 3B). Moreover, high-power views of DAT immunoreactivity revealed that abnormal foci were represented by clusters of rings (Fig. 3C and D) whose cores were positive for markers of the adjacent cell (Fig. 3D), indicating a tight interdigitation of these axonal structures with the neighboring cell.

**Fig 3.**
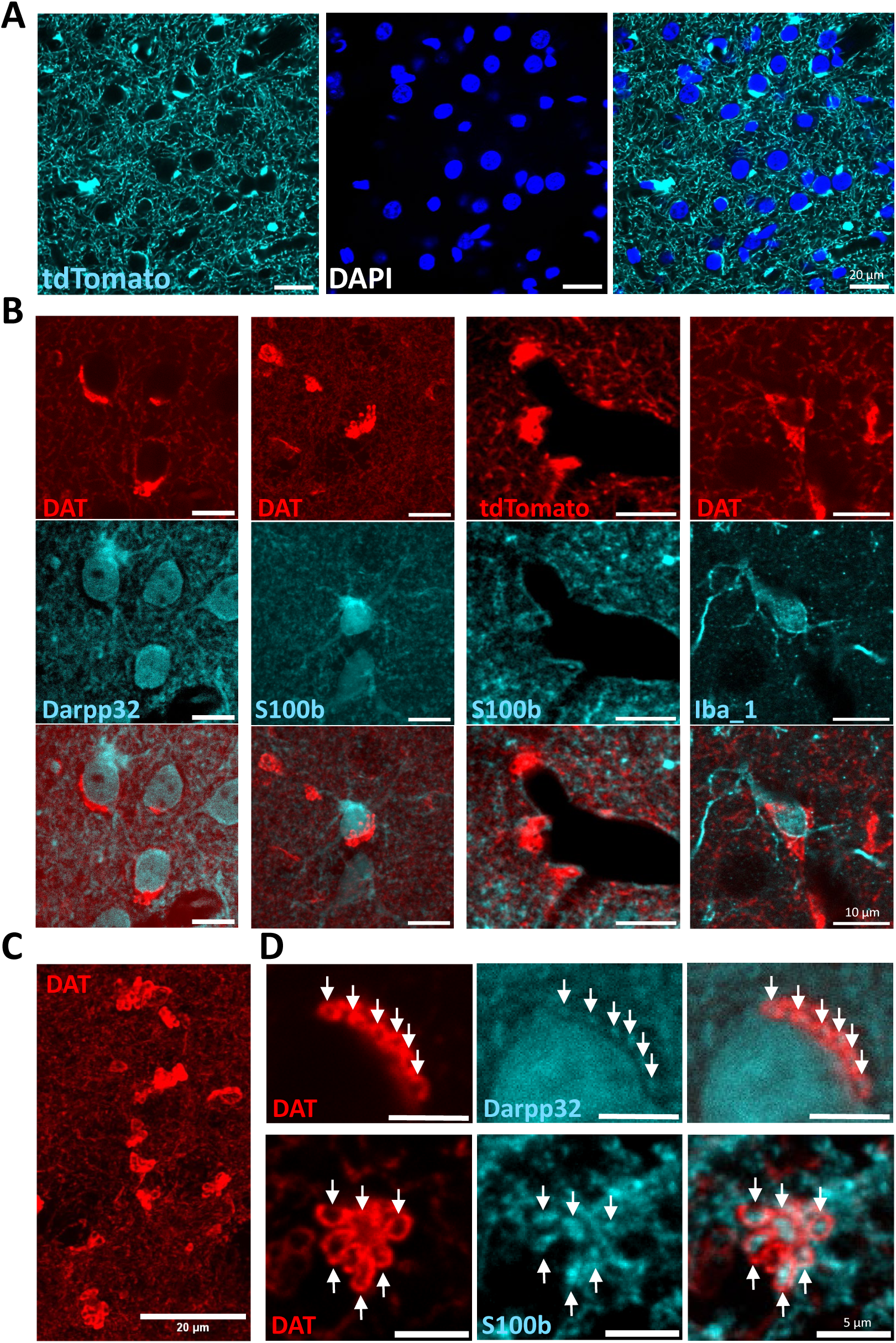
Dystrophic axon terminals of DAergic Synj1^RQ^ neurons are generally localized next to cell bodies and blood vessels. **A**. Juxtaposition of tdTomato-positive dystrophic axon terminals to DAPI-positive cell bodies. **B**. Double immunofluorescence for DAT and for markers of different cell populations: Darpp32 (marker of medium spiny neurons), S100b (marker of astrocytes), Iba1 (marker of microglia). Note that large DAT-poisitve clusters are generally adjacent to cell bodies of other cells. **C**. Abnormal foci of DAT immunoreactivity are represented by clusters of rings. **D**. Double immunofluorescence for DAT and for DARPP32 or for S100b shows that dystrophic axon contains evaginations of the neighboring cell. Scale bars: **A**, 20 µm; **B**, 10 µm; **C**, 20 µm; **D**, 5 µm. **A**: 3 month-old male; **B**-**D**: 3-5 month-old females and males.

Inspection at high resolution of the tdTomato fluorescence of 3D volumes of delipidated DLS revealed occasional dystrophic axons that made contact with multiple cell bodies (Fig. S2). However, generally, dystrophic axonal foci did not appear to be interconnected, suggesting that they reflect local changes rather than disruption of an entire axonal arbor.

### Dystrophic foci correspond to membrane whorls often surrounding evaginations of the adjacent cell

To unambiguously determine the ultrastructure of dystrophic axonal foci, we performed correlative light-electron microscopy (CLEM). In order to precisely identify the tdTomato fluorescent structures of interest, we used for EM a 3D technique: Focused Ion Beam Scanning Electron Microscopy (FIB-SEM) (30, 31). Vibratome sections (50 µm thick) were first observed by fluorescence microscopy to define regions of interest. Sections were then embedded in Epon, and Epon blocks were processed for FIB-SEM initially using a commercial Zeiss Crossbeam 550 machine. Three-domensional EM volumes were subsequently aligned with fluorescence images, revealing that foci of tdTomato fluorescence corresponded to clusters of membrane whorls or onion-like structures as illustrated in Fig. 4 and movies S4 and S5, which show two such elements close to a blood vessel. Within a cluster, these elements were interconnected, indicating that they originate from the same axon. This was also clearly exemplified by the 3D reconstruction of a very large branched mitochondrion in one of the foci shown in Fig. 4F, where such branches extended into different membrane whorls within the same cluster (Fig. 4K and L). The cores of these “onions” often harbored evaginations of nearby cells, consistent with light microscopy observations (asterisks in Fig. 4C-H).

**Fig. 4.**
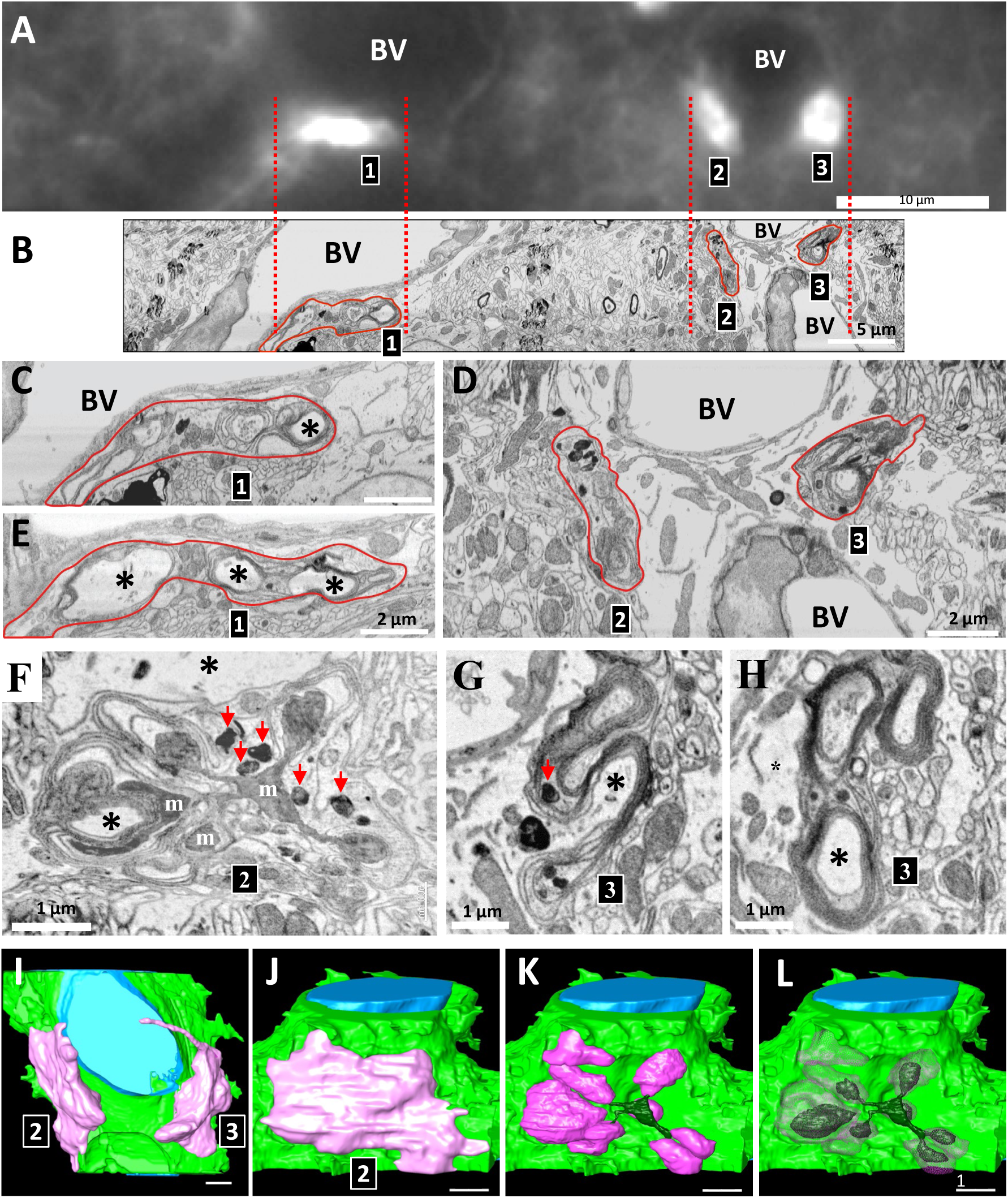
CLEM-FIB-SEM analysis of tdTomato expressing dystrophic DAergic axons in the striatum of a Synj1^RQ^ mouse (4 month-old female). **A** and **B**. tdTomato fluorescence (top) and FIB-SEM image of the same region of the striatum at the same magnification. The FIB-SEM image (**B**) shows that the three clusters (numbered 1,2 and 3) of tdTomato fluorescence visible in (**A**), correspond to multilayered membrane structures. **C** and **D**. Same FIB-SEM images as in **B** but at higher magnification. **E**. Another view of cluster 1 but at a different level in the 3D volume. **F**-**H**. Different views of clusters 2 and 3 at different levels within the 3D volume and at higher magnification relative to field **D**. Black asterisks in fields **C** and **E**-**H** indicate evaginations of astrocytes engulfed by dystrophic axons (see also movie 2). m = mitochondria. **I** and **J**. 3D surface view of the dystrophic axons forming clusters 2 and 3. Green = surface of an astrocyte; blue = blood vessel lumen; pink = surface view of the dystrophic terminal. **K** and **L**. Reconstruction of membranes within the dystrophic terminal. magenta = membrane whorls, dark green = mitochondria. The images of this figure were generated from movie S2 and S3. Scale bars: **A**, 10 µm; **B**, 5 µm; **C**-**E**, 2 µm; **F**-**L**, 1 µm.

### Membrane whorls derive from massive invaginations of the plasma membrane

Higher resolution views of Synj1^RQ^ mutant striata were obtained using a customized FIB-SEM machine at Janelia Research Campus, as shown in Fig. 5 (32, 33). The dystrophic axon shown in this figure comprises multiple whorls that enwrap evaginations of a neighboring neuron (Fig. 5B, D and E; movies S6 and S7).

**Fig. 5.**
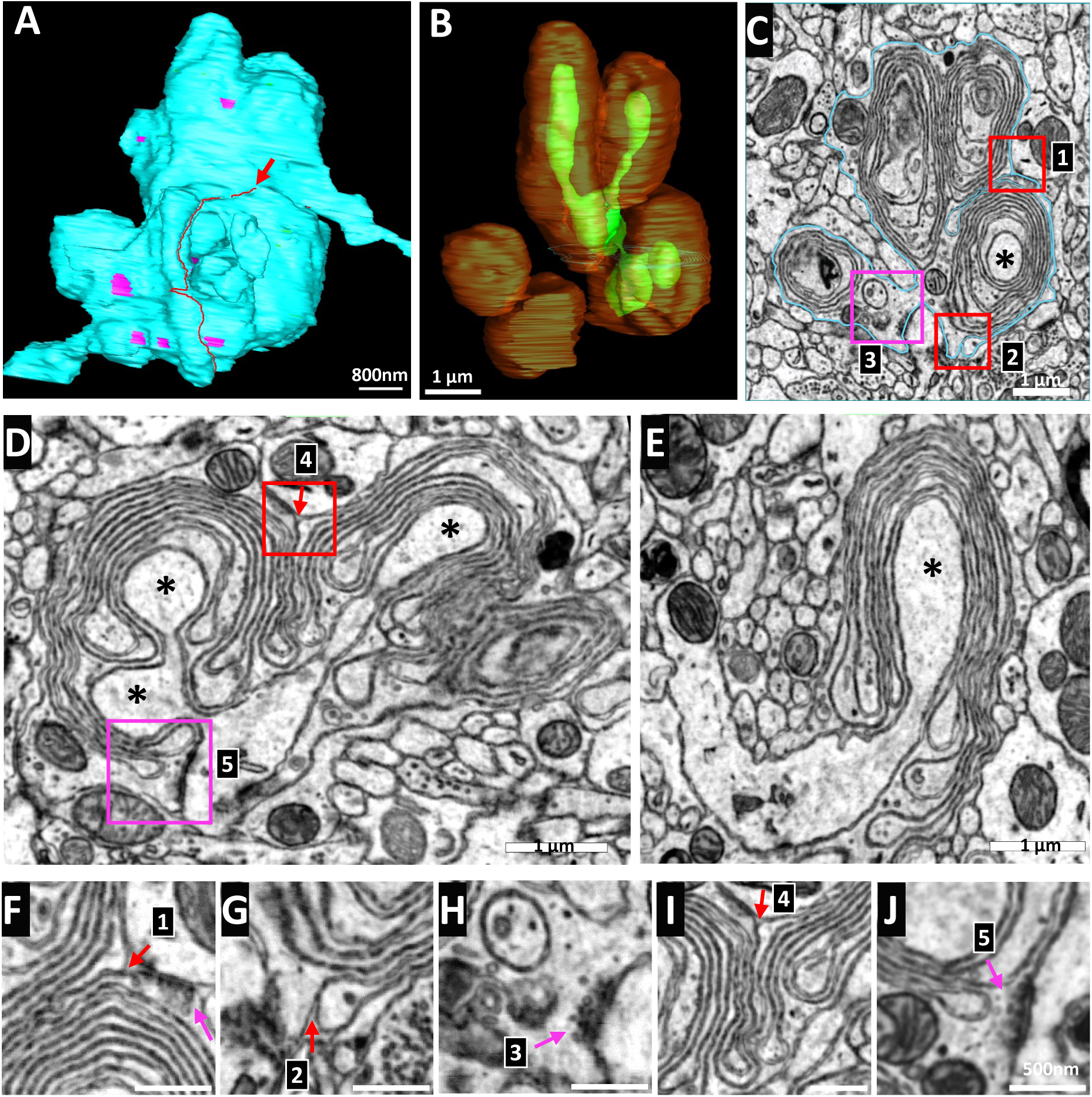
Analysis of a branched dystrophic nerve terminal of Synj1^RQ^ striatum with a customized high resolution FIB-SEM device (8 month-old female). **A** and **B**. 3D reconstruction of a dystrophic nerve terminal. **A** shows a surface view, with a red line (arrows) indicating where the outer plasma membrane folds inwards to generate the lamellar onion-like structures. **B** shows the onion-like structures (brown) which enclose evaginations (green) of an adjacent neuron. A synapse of the dystrophic terminal onto the neuron is visible in square #5 shown at high magnification in field J. **C** - **E** Single FIB-SEM slices of the 3D volume of the dystrophic nerve terminal shown in **A** and **B**. A blue line in **C** outlines the outer plasma membrane. Asterisks mark evaginations of an adjacent neuron. **F** - **J**. High magnification views of the numbered rectangles in **C** and **D**. Arrows in **F**, **G** and **I** point to where the outer cell plasma membrane invaginates to generate the massive onion-like whorls. Arrows in H and J point to two small synaptic vesicle clusters at active zones where the dystrophic nerve terminal makes synaptic contacts with a neighboring cell. Scale bars: **A**, 800 nm; **B**-**E**, 1 µm; **F**-**J**, 500 nm.

Close inspection of the multilayered membrane structures in FIB-SEM volumes (see arrows in fig. 5C, D, F, G and I) allowed us to identify them as sheet-like invaginations of the axonal plasma membrane that wrap around portions of the adjacent cell (Fig. 5D and E). While these images are reminiscent of myelin, in this case it is the plasma membrane of the DA axons that forms these invaginations, as discussed previously (19), and not the plasma membrane of oligodendrocytes. This was consistent with the intense immunoreactivity of membrane whorls for DAT (Fig. 1C, 3C and 3D), a marker of the axonal plasma membrane of DA neurons. The shape of these structures explains why large foci of DAT immunoreactivity have a ring-like shape when observed at high-power fluorescence microscopy (see Fig. 3C and 3D). Further confirming the axonal identity of these structures was the continuity with presynaptic active zones (Fig. 5F, H and J) and the presence of scattered synaptic vesicles among lamellae, as it can be seen more clearly in specimens analyzed by conventional TEM (Fig. 6).

**Fig. 6.**
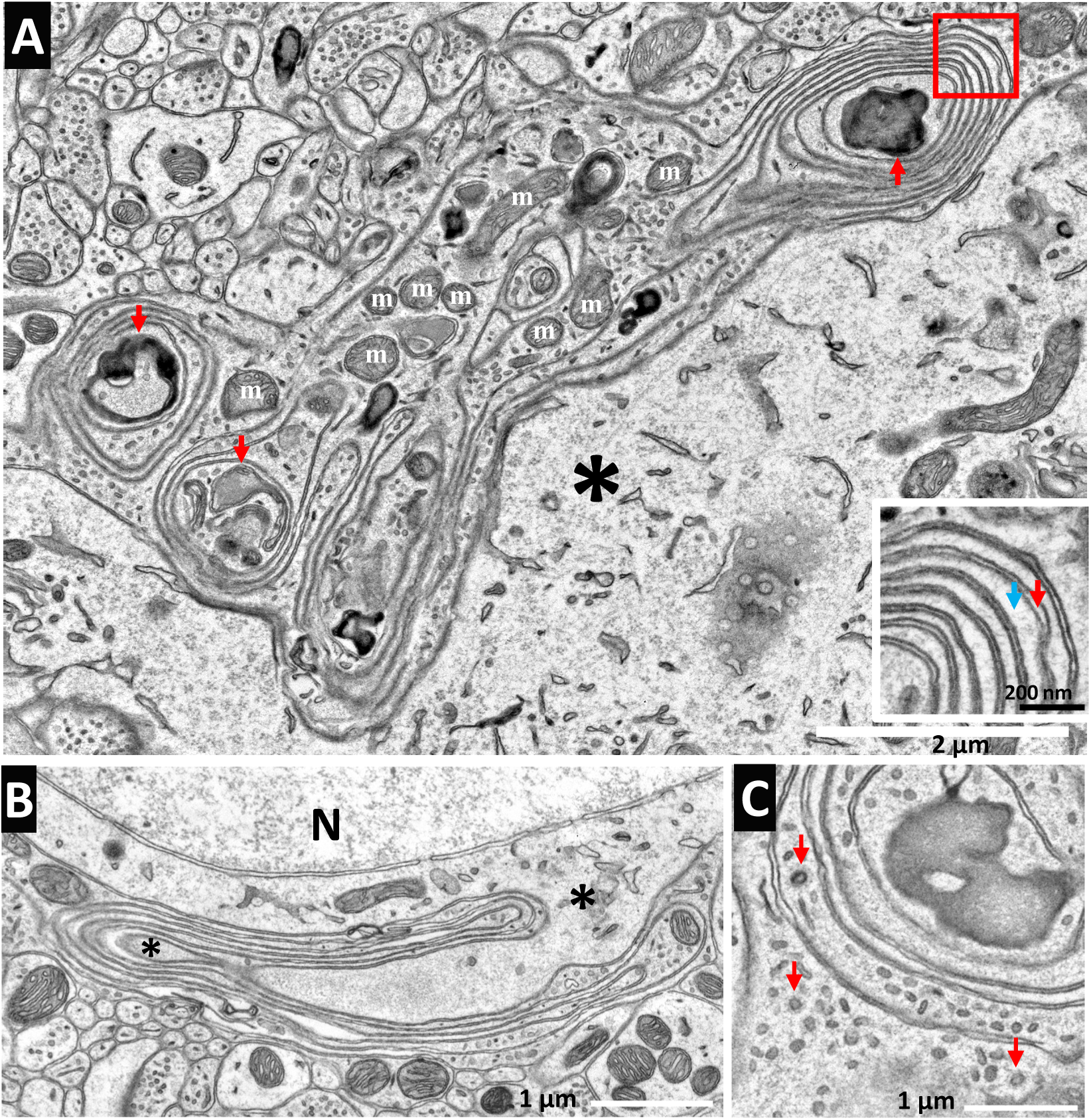
Conventional TEM demonstrating presence of trapped organelles in the membrane whorls (1.5 month-old male). **A**. Example of lamellar plasma membrane invaginations of a dystrophic axon including a variety of cell organelles among individual lamellae. Dense organelles (red arrows), possibly degenerating mitochondria or lysosomes, are present in the core of onion-like whorls. An asterisk indicates a neighboring cell. M = mitochondria; SV = synaptic vesicles. Inset: high magnification view of the region enclosed by a red square in the main field; red and blue arrows point to extracellular space and cytosolic space respectively. **B**. Multilayered plasma membrane invagination enclosing the evagination of a neighboring cell (asterisk). **C**. SVs and three clathrin coated vesicles are visible among lamellae. Scale bars: **A**, 2 µm, inset, 200 nm; **B** and **C**, 1 µm.

In some cases, the core of onion-like structures contained organelles with an electron-dense appearance (Fig. 4F and 6A). At least some of them appeared to be degenerating mitochondria, others may be lysosomes or autophagosomes. Possibly, organelles trapped in these structures, either in the dystrophic terminals, or in the evaginations of the cells that they enwrap, undergo degenerative changes due to their partial segregation from other portions of the cell.

### Synaptic vesicle markers of DAergic neurons are not enriched in the membrane whorls

As Synj1 is implicated in the endocytic recycling of synaptic vesicles (5, 8, 34, 35), it was of interest to determine whether vesicle proteins known to be enriched in synaptic vesicles of DAergic nerve terminals, SV2C (36, 37) and vMAT2 (38), were enriched in the whorls. SV2C and vMAT2 immunoreactivity were concentrated in abnormal axonal foci, but did not overlap with DAT, indicating that formation of the massive plasma membrane invagination is a process distinct from the endocytic events that recapture synaptic vesicle proteins after their exocytosis (Fig. 7A and B).

**Fig. 7.**
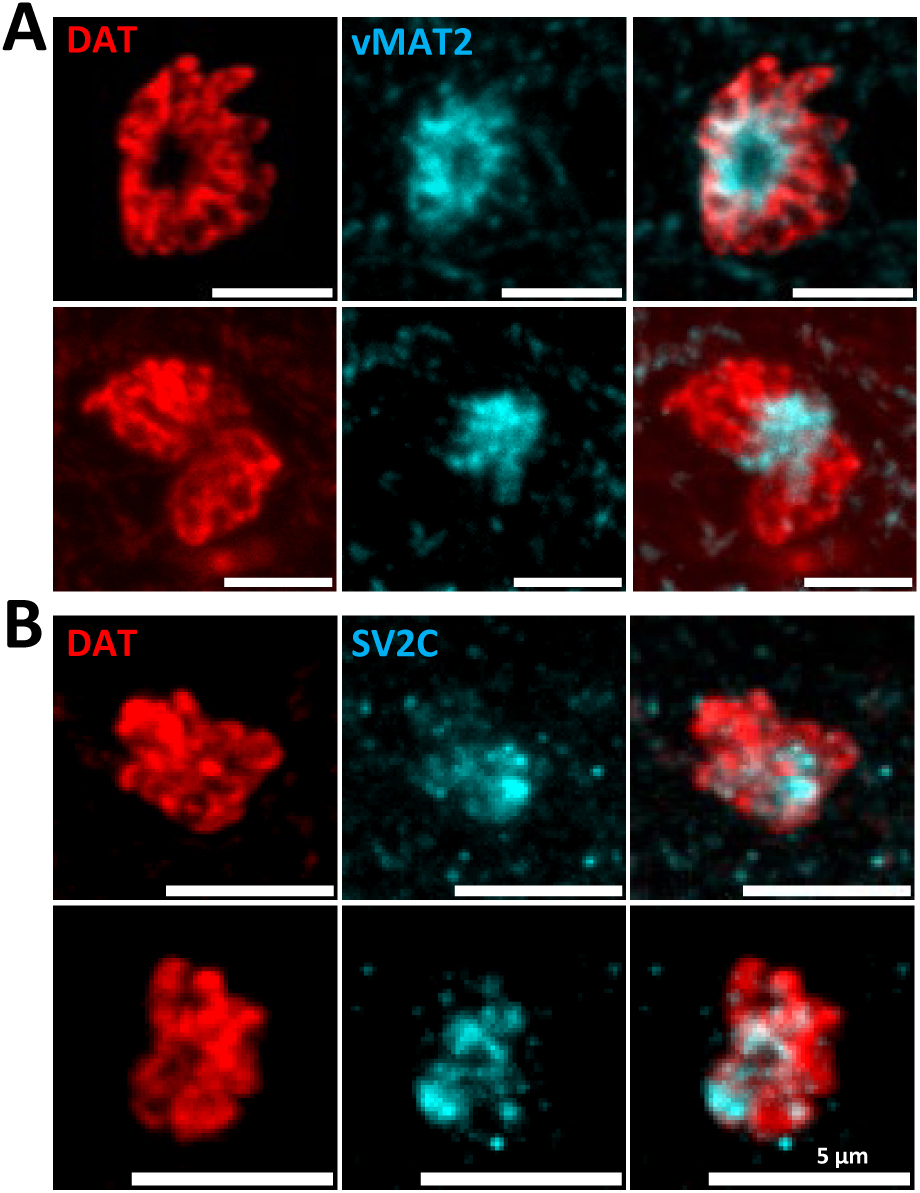
Segregation of DAT from markers of dopamine containing synaptic vesicle proteins in dystrophic nerve terminals. **A** and **B**. Double immunofluorescence for DAT and the synaptic vesicle membrane proteins vMAT2 and SV2C. The different localization of the two vesicular markers relative to DAT within the dystrophic axonal foci indicate that the membrane invaginations are not enriched in synaptic vesicle proteins. Scale bars: 5 µm. (4-5 month-old males and females).

## DISCUSSION

Animal models of disease are powerful experimental tools to dissect disease mechanisms. While more than 20 mutations responsible for familial forms of PD have already been identified (1, 3), in most cases the introduction of the corresponding mutations in mice does not result in an obvious disease phenotype. Mice with the R258Q patient mutation in Synj1 have at least some of the neurological phenotypes observed in human patients (19, 20). Importantly, dystrophic changes in a subpopulation of DAergic axons are present in the striata of these mice and these changes are largely restricted to the dorsolateral striatal regions known to be primarily involved in the motor dysfunction of PD patients (39–41). As we have shown, these changes are accompanied by a defect in evoked DA release specifically in the same striatal region. Thus, studies of these mice may help shed light on the special vulnerability of these neurons.

Here we have further elucidated the nature of these focal dystrophic changes. Using FIB-SEM CLEM, we have proven that they are represented by massive invaginations of the axonal plasma membrane that form onion-like structures somewhat reminiscent of myelin. However, besides the difference in the cell type involved (neurons rather than oligodendrocytes), the whorls represent invaginations, rather than evaginations of the plasma membrane as in myelin. Moreover, in contrast to the layers of myelin, where membranes are tightly apposed with no intervening cytoplasm, lamellae of these axonal invaginations contain thin layers of cytoplasm and in some cases also small organelles. As dystrophic segments are observed only in a minor subpopulation of DA axons even in striatal regions where they are most abundant, it remains unclear whether many axons of few neurons, or few axonal branches of different neurons, are affected. We have further shown that these onion-like structures trap evaginations of adjacent cell bodies, raising the possibility of non-cell autonomous effects in the pathological manifestations. Much remains to be learned about mechanisms underlying these striking pathological changes.

DAT is a major component of these whorls. In contrast, synaptic vesicle proteins known to be enriched in vesicles of DA nerve terminals, such as SV2C and vMAT2 (36–38), do not colocalize with DAT-positive membrane whorls although they are enriched in these dystrophic axonal segments. Possibly they reflect synaptic vesicles trapped in the axon dilations. This speaks against the possibility that the dramatic expansion of the plasma membrane responsible for the formation of the membrane invaginations may simply reflect synaptic vesicle exocytosis not balanced by synaptic vesicle endocytosis. In fact, exocytosis of synaptic vesicles may be impaired in mutant terminals, as supported by the defect in evoked DA release detected by our functional experiments. A selective stranding at the plasma membrane of DAT delivered by some form of constitutive exocytosis, must be involved.

Patients with loss-of-function mutations in auxilin (alias: DNAJC6 or PARK19) develop early onset parkinsonism with clinical manifestations very similar to those of patients with the SYNJ1^RQ^ mutation (15, 16, 42, 43). Interestingly, dystrophic changes in DAergic axons very similar - in morphology, localization and enrichment in DAT - to those observed in the DLS of Synj1^RQ^ are present in the DLS of auxilin KO mice (20, 44). Auxilin and Synj1 are functional partners in endocytosis (45). The PI(4,5)P_2_ phosphatase activity of Synj1 allows release of clathrin adaptors and other endocytic factors anchored to the membrane via an interaction with PI(4,5)P_2_ (5, 8, 9). Auxilin, a cofactor of the ATPase HSC70, is implicated in clathrin coat disassembly (46, 47). In agreement with the functional partnership of these two proteins with house-keeping functions in neurons, a decreased number of synaptic vesicles and an increased number of clathrin-coated vesicles are observed at synapses throughout the brain, including at striatal synapses, of mice harboring Synj1 or auxilin loss-of-function mutations (5, 19, 20, 44, 48). The changes observed in dystrophic DAergic neurons, however, do not represent an exaggeration of this phenotype. In fact, only occasional clathrin coated vesicles are observed in proximity to the membrane whorls. Perhaps, as mentioned above, the onset of the dystrophic changes correlates with a shutdown of SV exocytosis in these terminals.

Recently, it was shown that abnormal dystrophic swelling of DAergic axons occurs also in the VS of mice in which the Synj1^RQ^ mutation is combined with the loss of auxilin (20), or in mice with conditional KO of Synj1 in DA neurons rather than simply harboring the Synj1^RQ^ mutation which selectively impairs the Sac1 phosphatase activity of this protein (24). Thus, it would appear that the axonal arbors of DA neurons are particularly vulnerable to the lack of Synj1 and auxilin function, with more severe vulnerability of the subpopulation projecting to the DLS.

It was reported that Synj1 haploinsufficiency results in increased levels of DAT in DAergic neurons, but in reduced surface expression of DAT (49). This would seem in contrast with our evidence that in mice with impaired Synj1 function a massive amount of DAT is trapped in the dramatically expanded plasma membrane of a subset of axons. However, while the Synj1^RQ^ mutation abolishes its 4-phosphatase activity (15, 16), reduced surface expression of DAT in mice with a *Synj1* KO allele was attributed to impaired 5-phosphatase activity (49). The trapping of DAT at the plasma membrane is also supported by the reports that in auxilin mutant mice, where similar DAT-enriched structures are observed, such structures are accessible to a membrane-impermeant DAT ligand (44).

A key open question is why the dystrophic changes described here in Synj1^RQ^ mice occur so specifically in a subpopulation of DAergic axons of the DLS. Our functional studies of evoked DA release in striatal slices have revealed a defect of such release in the same brain region and not in the VS. It is unlikely, however, that such a defect may be accounted for by the relatively small percentage of dystrophic axons. We favor a model in which the Synj1^RQ^ mutation results in a general selective impairment of DA neurons projecting to the dorsal striatum. In a subset of axonal segments, this functional impairment may reach a threshold beyond which the normal balance between exocytosis and endocytic mechanisms, including constitutive exocytosis-endocytosis to maintain plasma membrane homeostasis, may be chronically perturbed, leading to massive plasma membrane invaginations and synaptic vesicle exocytosis shut down. Elucidating the precise causes for the selective vulnerability of DAergic neurons to the lack of Synj1 and auxilin, and more so of the subset of neurons projecting to the DLS, remains an important priority for future studies.

## Acknowledgments

We thank Xinran Liu and Morven Graham for assistance in FIB-SEM imaging carried out in the CCMI facility at Yale and Yulan (Mumu) Fang for technical assistance in TEM. This work was supported in part by grants to P.D.C. from the NIH (DA018343), the Parkinson’s Foundation (PF-RCE-1946), the Freedom Together Foundation (2024-4715 to DJS), and the Aligning Science Across Parkinson’s through the Michael J. Fox Foundation for Parkinson’s Research (ASAP-000580 to P.D.C.; ASAP-020551 to D.J.S.). For the purpose of open access, the authors have applied a CC BY 4.0 public copyright license.

## Author contributions

PDC, YW conceived the project and wrote the manuscript with contributions from DJS. YW carried out most of the experiments and analyses, including the FIBSEM performed at the Yale CCMI, PX carried out the mouse breedings to generate DAT^IRES^cre; Ai9; WT or *Synj1*^RQ/RQ^ mice and to validate them, JM under the supervision of DJS carried out the DA release assays, CSX, KJH and HFH generated the FIBSEM dataset at Janelia Research Campus, LS helped in studies of delipidated mouse brains, MC generated the Synj1^RQ^ brain tissue used for FIB-SEM analysis carried out at Janelia Research Campus. All authors read and approved the final manuscript.

## Competing interest

All authors declare no financial competing interests. Dr. Robert Edwards (a reviewer of this manuscript) and Dr. James Surmeier (an author of this manuscript) were coauthors in a 2024 review (DOI: 10.1002/mds.29897).

## METHODS

### Detailed methods are provided in the Supporting Information

#### Brain Immunofluorescence

Mice were anesthetized with Ketamine/Xylazine and transcardially perfused with 4% PFA/0.05% Glutaraldehyde in 0.1M PB (pH 7.4). Brains were post-fixed overnight at 4°C and sectioned at 25–30 µm with a vibratome. Floating sections were incubated with primary antibodies followed by Alexa-conjugated secondary antibodies and mounted with ProLong Gold. Images were acquired on an Olympus VS200 (20×) or Nikon Ti2-E/CSU-W1 SoRa (60×) and analyzed with Fiji.

#### Brain tissue clearing and imaging

Brain were cleared using the SHIELD protocol (LifeCanvas). Perfused brains were post-fixed overnight, crosslinked in SHIELD OFF/ON solutions, delipidated (1 week, 45°C), and cleared in EasyIndex (RI = 1.52). Light-sheet imaging was performed on a SmartSPIM light-sheet microscope (LifeCanvas, 9×/0.3 NA). A 4×6 tiled dataset was acquired, stitched, and the striatum segmented based on tdTomato fluorescence. Dystrophic axonal segments (large puncta of tdTomato fluorescence) were identified by threshold-based segmentation of tdTomato-positive puncta using Zeiss Arivis Pro. For confocal imaging, 100 µm sagittal sections were re-cleared in EasyIndex and imaged on a Nikon Ti2-E/CSU-W1 SoRa (60× oil-immersion objective). 3D stacks (200×200×100 µm³, 108 nm XY/100 nm z) were analyzed with Imaris and Dragonfly Pro.

#### FIB-SEM CLEM at the Yale CCMI facility

Mice were perfused and fixed as described above. Sagittal vibratome sections (50 µm) were imaged on an Olympus VS200 (40×) slide scanner. Selected sections were then postfixed in glutaraldehyde and processed through the OTO post-fixation staining protocol (see detailed methods) and Epon embedded. After platinum sputter-coating and SEM imaging, regions of interest were identified by correlating tissue landmarks with tdTomato clusters and processed (FIB milling and SEM imaging) through a Zeiss Crossbeam 550 FIB-SEM at 8 nm/voxel. 3D volumes were reconstructed with Dragonfly Pro.

#### FIBSEM at HHMI Janelia Research Campus

Mouse perfusion and epon embedding of brain smaples were performed as described in detailed methods. Epon-embedded samples were trimmed, re-embedded in durcupan, and coated with 10-nm gold/100-nm carbon. FIB-SEM imaging was carried out with a custom FEI Magnum FIB/Zeiss Merlin SEM (200-pA, 700 V electron beam; 7-nA gallium ion milling) at 8-nm isotropic voxels with backscattered electron detection. 3D reconstructions were generated by semi-manual membrane tracing in 3dmod.

#### Conventional TEM imaging

Striatal sections prepared as described above for FIBSEM (YALE CCMI facility) were cut with a Leica ultramicrotome to generate 60 nm ultrathin sections. Sections were post-stained with uranyl acetate substitute (Uranyless, EMS), followed by a lead citrate (EMS) solution. Sections were observed in a Talos L 120 C TEM microscope at 80 kV and images were taken with Velox software and a 4k × 4 K Ceta CMOS Camera (Thermo Fisher Scientific).

#### Ex vivo slice preparation for the analysis of DA release

Mice were anesthetized, perfused with ice-cold slicing aCSF (containing 49.14 mM NaCl, 2.5 mM KCl, 1.43 mM NaH2PO4, 25 mM NaHCO3, 25 mM glucose, 99.32 mM sucrose, 10 mM MgCl2, and 0.5 mM CaCl2), and decapitated. Coronal slices (220 µm) were cut with a Leica VT1200S vibratome and recovered in recording aCSF (containing 135.7 mM NaCl, 2.5 mM KCl, 1.25 mM NaH2PO4, 25 mM NaHCO3, 2 mM CaCl2, 1 mM MgCl2, and 3.5 mM glucose) (30 min, 34°C; then ≥15 min, RT). All solutions were pH 7.4, ∼310 mOsm, 95% O₂/5% CO₂.

#### Two-photon laser scanning microscope (2PLSM) workstation

Two-photon imaging used a Bruker Ultima system (Olympus BX-51WIF, 60×/1.0 NA) with a Coherent Chameleon Ti:sapphire laser (690–1040 nm), Pockels’ cell power control, and non-descanned PMT detection in green (490–560 nm) and red (580–630 nm) channels. Images were acquired at 0.393 µm/pixel, 8 µs dwell time, 39.2 fps.

#### DA release measurements

Synj1^RQ^ or control mice received AAV5-CAG-dLight1.3b (500 nl) into DLS or NAc at P30; experiments were performed at P45. Brain slices were prepared as above. dLight was imaged at 920 nm (2PLSM, 20 µm spiral scan, 39.2 fps) with DA release evoked by electrical stimulation (600 µA, 0.2 ms, 1 or 5 pulses at 10 Hz) via a tungsten in the presence of 10µM mecamylamine, 10µM sulpiride, and 10µM gabazine. Signals are expressed as %ΔF max (calibrated with 100 µM DA) or Δf/f0 normalized to the first response.

#### Analysis and statistics

Images were analyzed in FIJI (50). Traces were processed in Excel (Microsoft) and GraphPad Prism8 (GraphPad software). Sample *n* represents the number of brain slices collected from *N* animals. Statistical significance (P < 0.05) was assessed using the Mann-Whitney U test for DA release comparisons and two-way repeated measures ANOVA with Šídák correction for vesicle recycling experiments.

## Data availability

The data, protocols, and key lab materials used and generated in this study are listed in a Key Resource Table alongside their persistent identifiers at [https://doi.org/10.5281/zenodo.22662279]. Analysis scripts is available at [Biostudies/S-BIAD3429]; all data cleaning, preprocessing, analysis, and visualization was performed using ImageJ, Zeiss Arivis Pro, Dragonfly Pro (Comet Technologies Canada Inc), Imaris, OLYMPUS OlyVIA 4.2, Atlas 5 (Fibics Incorporated), Velox (Thermo Fisher Scientific), 3dmod, Handbreak, Excel and GraphPad Prism8.

## Notes

### Competing Interest Statement

The authors have declared no competing interest.

### Summary of Updates

This is an updated preprint reflecting feedback from peer reviewers. Added Fig. S2: Analysis of dLight signal ΔF/F max and decay across genotypes. Updated the Methods section on dopamine release measurement. Corrected typos and grammatical errors throughout the text.

## REFERENCES

1. G. Cogan, S. Lesage, A. Brice, The genetics of autosomal recessive early-onset Parkinson’s disease. Curr Opin Neurobiol 95, 103141 (2025).

2. I. Boura et al., The genetic architecture of Parkinson’s disease on the Island of Crete. NPJ Parkinsons Dis 11, 354 (2025).

3. C. Blauwendraat, M. A. Nalls, A. B. Singleton, The genetic architecture of Parkinson’s disease. Lancet Neurol 19, 170–178 (2020).

4. P. S. McPherson et al., A presynaptic inositol-5-phosphatase. Nature 379, 353–357 (1996).

5. O. Cremona et al., Essential role of phosphoinositide metabolism in synaptic vesicle recycling. Cell 99, 179–188 (1999).

6. P. De Camilli, Roles in physiology and disease of the inositol phosphatase synaptojanin 1. Biochim Biophys Acta Mol Cell Biol Lipids 1871, 159708 (2026).

7. M. R. Wenk, P. De Camilli, Protein-lipid interactions and phosphoinositide metabolism in membrane traffic: insights from vesicle recycling in nerve terminals. Proc Natl Acad Sci U S A 101, 8262–8269 (2004).

8. I. Milosevic et al., Recruitment of endophilin to clathrin-coated pit necks is required for efficient vesicle uncoating after fission. Neuron 72, 587–601 (2011).

9. P. Verstreken et al., Synaptojanin is recruited by endophilin to promote synaptic vesicle uncoating. Neuron 40, 733–748 (2003).

10. H. Choudhry, M. Aggarwal, P. Y. Pan, Mini-review: Synaptojanin 1 and its implications in membrane trafficking. Neurosci Lett 765, 136288 (2021).

11. L. W. Gong, P. De Camilli, Regulation of postsynaptic AMPA responses by synaptojanin 1. Proc Natl Acad Sci U S A 105, 17561–17566 (2008).

12. S. Yang et al., Presynaptic autophagy is coupled to the synaptic vesicle cycle via ATG-9. Neuron 110, 824–840.e810 (2022).

13. K. Hardies et al., Loss of SYNJ1 dual phosphatase activity leads to early onset refractory seizures and progressive neurological decline. Brain 139, 2420–2430 (2016).

14. D. A. Dyment et al., Homozygous nonsense mutation in SYNJ1 associated with intractable epilepsy and tau pathology. Neurobiol Aging 36, 1222.e1221–1225 (2015).

15. C. E. Krebs et al., The Sac1 domain of SYNJ1 identified mutated in a family with early-onset progressive Parkinsonism with generalized seizures. Hum Mutat 34, 1200–1207 (2013).

16. M. Quadri et al., Mutation in the SYNJ1 gene associated with autosomal recessive, early-onset Parkinsonism. Hum Mutat 34, 1208–1215 (2013).

17. K. Senkevich et al., Are rare heterozygous SYNJ1 variants associated with Parkinson’s disease? NPJ Parkinsons Dis 10, 201 (2024).

18. S. Lesage et al., Clinical Variability of SYNJ1-Associated Early-Onset Parkinsonism. Front Neurol 12, 648457 (2021).

19. M. Cao et al., Parkinson Sac Domain Mutation in Synaptojanin 1 Impairs Clathrin Uncoating at Synapses and Triggers Dystrophic Changes in Dopaminergic Axons. Neuron 93, 882–896.e885 (2017).

20. X. Y. Ng et al., Mutations in Parkinsonism-linked endocytic proteins synaptojanin1 and auxilin have synergistic effects on dopaminergic axonal pathology. NPJ Parkinsons Dis 9, 26 (2023).

21. M. Cao, D. Park, Y. Wu, P. De Camilli, Absence of Sac2/INPP5F enhances the phenotype of a Parkinson’s disease mutation of synaptojanin 1. Proc Natl Acad Sci U S A 117, 12428–12434 (2020).

22. L. Madisen et al., A robust and high-throughput Cre reporting and characterization system for the whole mouse brain. Nat Neurosci 13, 133–140 (2010).

23. C. M. Bäckman et al., Characterization of a mouse strain expressing Cre recombinase from the 3’ untranslated region of the dopamine transporter locus. Genesis 44, 383–390 (2006).

24. Y. Lin et al., Loss of Synaptojanin 1 in dopamine neurons triggers synaptic degeneration and striatal TH interneuron compensation. bioRxiv 10.1101/2025.05.14.653914, 2025.2005.2014.653914 (2026).

25. Y. G. Park et al., Protection of tissue physicochemical properties using polyfunctional crosslinkers. Nat Biotechnol 10.1038/nbt.4281 (2018).

26. A. Nishi, G. L. Snyder, P. Greengard, Bidirectional regulation of DARPP-32 phosphorylation by dopamine. J Neurosci 17, 8147–8155 (1997).

27. S. K. Ludwin, J. C. Kosek, L. F. Eng, The topographical distribution of S-100 and GFA proteins in the adult rat brain: an immunohistochemical study using horseradish peroxidase-labelled antibodies. J Comp Neurol 165, 197–207 (1976).

28. J. Du et al., S100B is selectively expressed by gray matter protoplasmic astrocytes and myelinating oligodendrocytes in the developing CNS. Mol Brain 14, 154 (2021).

29. Y. Imai, I. Ibata, D. Ito, K. Ohsawa, S. Kohsaka, A novel gene iba1 in the major histocompatibility complex class III region encoding an EF hand protein expressed in a monocytic lineage. Biochem Biophys Res Commun 224, 855–862 (1996).

30. Y. Wu et al., Contacts between the endoplasmic reticulum and other membranes in neurons. Proc Natl Acad Sci U S A 114, E4859–e4867 (2017).

31. C. S. Xu et al., An open-access volume electron microscopy atlas of whole cells and tissues. Nature 599, 147–151 (2021).

32. C. S. Xu et al., Enhanced FIB-SEM systems for large-volume 3D imaging. Elife 6 (2017).

33. S. Pang, C. S. Xu, Methods of enhanced FIB-SEM sample preparation and image acquisition. Methods Cell Biol 177, 269–300 (2023).

34. M. Mani et al., The dual phosphatase activity of synaptojanin1 is required for both efficient synaptic vesicle endocytosis and reavailability at nerve terminals. Neuron 56, 1004–1018 (2007).

35. W. T. Kim et al., Delayed reentry of recycling vesicles into the fusion-competent synaptic vesicle pool in synaptojanin 1 knockout mice. Proc Natl Acad Sci U S A 99, 17143–17148 (2002).

36. R. Janz, T. C. Südhof, SV2C is a synaptic vesicle protein with an unusually restricted localization: anatomy of a synaptic vesicle protein family. Neuroscience 94, 1279–1290 (1999).

37. A. R. Dunn et al., Synaptic vesicle glycoprotein 2C (SV2C) modulates dopamine release and is disrupted in Parkinson disease. Proc Natl Acad Sci U S A 114, E2253–e2262 (2017).

38. B. Onoa, H. Li, J. A. Gagnon-Bartsch, L. A. Elias, R. H. Edwards, Vesicular monoamine and glutamate transporters select distinct synaptic vesicle recycling pathways. J Neurosci 30, 7917–7927 (2010).

39. A. Fushiki et al., A Vulnerable Subtype of Dopaminergic Neurons Drives Early Motor Deficits in Parkinson’s Disease. bioRxiv 10.1101/2024.12.20.629776 (2024).

40. I. Mantas et al., Anxa1+ dopamine neuron vulnerability defines prodromal Parkinson’s disease bradykinesia and procedural motor learning impairment. bioRxiv 10.1101/2024.12.22.629963, 2024.2012.2022.629963 (2025).

41. M. R. Hilário, R. M. Costa, High on habits. Front Neurosci 2, 208–217 (2008).

42. S. Edvardson et al., A deleterious mutation in DNAJC6 encoding the neuronal-specific clathrin-uncoating co-chaperone auxilin, is associated with juvenile parkinsonism. PLoS One 7, e36458 (2012).

43. Ç. Köroğlu, L. Baysal, M. Cetinkaya, H. Karasoy, A. Tolun, DNAJC6 is responsible for juvenile parkinsonism with phenotypic variability. Parkinsonism Relat Disord 19, 320–324 (2013).

44. D. J. Vidyadhara et al., Dopamine transporter and synaptic vesicle sorting defects underlie auxilin-associated Parkinson’s disease. Cell Rep 42, 112231 (2023).

45. Y. Saheki, P. De Camilli, Synaptic vesicle endocytosis. Cold Spring Harb Perspect Biol 4, a005645 (2012).

46. K. He et al., Dynamics of Auxilin 1 and GAK in clathrin-mediated traffic. J Cell Biol 219 (2020).

47. E. Eisenberg, L. E. Greene, Multiple roles of auxilin and hsc70 in clathrin-mediated endocytosis. Traffic 8, 640–646 (2007).

48. Y. I. Yim et al., Endocytosis and clathrin-uncoating defects at synapses of auxilin knockout mice. Proc Natl Acad Sci U S A 107, 4412–4417 (2010).

49. J. Saenz et al., Parkinson’s disease gene, Synaptojanin1, dysregulates the surface maintenance of the dopamine transporter. NPJ Parkinsons Dis 10, 148 (2024).

50. J. Schindelin et al., Fiji: an open-source platform for biological-image analysis. Nat Methods 9, 676–682 (2012).

