## Supplemental Materials for "Dystrophic changes of nigrostriatal axons harboring a Synj1 Parkinson mutation suggest catastrophic failure of endocytic mechanisms"

### Extended Methods

**Antibodies.** Rabbit anti-DARPP-32 (2306, RRID: AB_823479) from Cell Signaling Technology; rat anti-DAT (MAB369, RRID: AB_2190413) from Millipore;; rabbit anti-SV2C (119203, RRID: AB_993023), from Synaptic Systems; mouse anti-S100b (SAB4200671, RRID: AB_3712829) from Sigma; rabbit anti-Iba1 (01919741, RRID: 839504) from FUJIFILM Wako Pure Chemical Corporation; rabbit anti-vMAT2 (MSFR107570, RRID: AB_2571857) from FRONTIER INSTITUE CO., LTD.

**Mice.** The following mouse strains were interbred to generate mice expressing tdTomato in DAergic neurons and harboring either the WT or R259Q *Synj1* allele: DAT$^{IREScre}$ (B6.SJL-Slc6a3$^{tm1.1(cre)Bkmn}$/J; RRID: IMSR_JAX-006660); Ai9 mice (B6.Cg-Gt(ROSA)26Sor$^{tm9(CAG-tdTomato)Hze}$/J; RRID: IMSR_JAX-007909); Synj1$^{RQ}$ mice in C57BL6 (C57BL/6-*Synj1*<R259Q>); RRID: MGI:6120537 (1). Mice heterozygous for DAT$^{IREScre}$ and either homozygous or heterozygous for the conditional tdTomato cassette were used for our experiments. All mice were maintained on a 12 h light/dark cycle with standard mouse chow and water ad libitum. All research and animal care procedures were approved by the Yale University Institutional Animal Care and Use Committee. A detailed procedure is available at: https://dx.doi.org/10.17504/protocols.io.36wgq23nygk5/v1.

**Brain Immunofluorescence.** Mice were anesthetized with a Ketamine (VetOne, NDC 13985-584-10)/Xylazine (Anased, NDC 59399-110-20) anesthetic cocktail injection, perfused transcardially with 37 °C pre-warmed 4% PFA and 0.05% Glutaraldehyde (Electron Microscopy Sciences) in 0.1M phosphate buffer (PB), PH7.4 and the brains were kept in the same fixative overnight at 4 °C. Brains were then washed 3 times for 20 min in 0.1M PB buffer and 25-30 µm thick coronal or sagittal sections were cut with a vibratome. Floating sections were incubated in 2% BSA, either 2% donkey (EMD Millipore, S30-100ML) or goat (ThermoFisher Scientific, 16210064) serum, 0.4% Triton X-100, 0.05% Tween20, 0.01% $NaN_3$, and 50mM $NH_4Cl$ in 0.1M PB buffer for 2 h at room temperature and then incubated with primary antibodies (diluted in the same buffer) overnight at 4 °C. Subsequently, sections were washed 3 times for 10 min with 0.1M PB buffer, then incubated with Alexa-conjugated secondary antibodies for 2 h at room temperature. Finally, sections were mounted with ProLong$^{TM}$Gold antifade reagent (Invitrogen, P36934) and sealed with nail polish. Images were acquired either with an Olympus slice view VS200 slide scanner equipped with a Hamamatsu Orca-Fusion camera and a 20x Olympus UPlanXApo objective or a Nikon Ti2-E inverted microscope (Yokogawa CSU-W1 SoRa, Nikon) equipped with a 60x SR Plan Apo IR oil-immersion objective. Images analysis was done with Fiji software. A detailed protocol is found at DOI: https://dx.doi.org/10.17504/protocols.io.14egn515zg5d/v1.

**Brain tissue clearing and imaging**. Brain tissue clearing was performed following the standard SHIELD protocol and passive delipidation procedure (LifeCanvas). Briefly, mice were perfused with 4% paraformaldehyde (PFA) and 0.05% glutaraldehyde in 0.1 M phosphate buffer (PB) as described above, and brains were post-fixed overnight in the same fixative at 4 °C. Tissues were then crosslinked by incubation in 20 mL SHIELD OFF solution for 3 days at 4 °C, followed by 20 mL SHIELD ON solution for 24 h at 37 °C with shaking (40 rpm, orbital shaker). Subsequently, brains were transferred to 20 mL delipidation buffer for 1 week at 45 °C with shaking (60 rpm). After delipidation, tissues were incubated under shaking (60 rpm) in 20 mL of 50% EasyIndex solution (RI = 1.52) in $H_2O$ for 24 h at 37 °C, followed by 100% EasyIndex at 37 °C for an additional 24 h until fully transparent.

*Full brain imaging by a light-sheet microscope.* The cleared mouse brain was embedded in an index-matching gel block and immersed in an imaging chamber filled with immersion oil (refractive index (RI) 1.52). Imaging was performed using a SmartSPIM light-sheet microscope (LifeCanvas Technologies) equipped with a 9×, 0.3 NA clearing objective (Applied Scientific Instrumentation). A total of 4 × 6 tiles were acquired and stitched together using the manufacturer's post-acquisition software. The left striatum was segmented and the neighboring areas were masked out based on tdTomato fluorescence intensity. The striatum fluorescence dataset was then preprocessed to remove background and nonspecific signal, followed by threshold-based segmentation of tdTomato-positive puncta corresponding to dystrophic dopaminergic axons using Zeiss Arivis Pro.

*Confocal imaging of vibratome sections*. For high-resolution confocal imaging, brains were washed in 0.1 M PB for 6 h with one buffer change. Sagittal brain sections (100 µm thick) were cut using a vibratome and re-incubated in EasyIndex for at least 20 min until transparency was restored. Sections were mounted on glass slides in one drop of EasyIndex Matched Immersion Oil (RI = 1.52), cover-slipped (12 mm), and imaged using a Nikon Ti2-E equipped with a Yokogawa CSU-W1 SoRa (Nikon) and a 60× SR Plan Apo IR oil-immersion objective (IMMERSION OIL TYPE F, RI = 1.518). 3D image stacks (200 × 200 × 100 $\mu m^3$) were acquired at high spatial resolution (108 × 108 nm per pixel in XY, 100 nm z-step), enabling detailed visualization of fine structures. Image datasets were subsequently processed and analyzed using Imaris (Oxford Instruments) and Dragonfly Pro (Comet Technologies Canada Inc.) software.

Details of the protocol can be found at:
https://sites.google.com/lifecanvastech.com/protocol/outline

**FIB-SEM CLEM at the Yale CCMI facility**. Mouse anesthesia and transcardial perfusion with fixative (4% paraformaldehyde and 0.05% glutaraldehyde in 0.1M PB buffer), followed by an overnight incubation with the same fixative and subsequent washes, were performed as described above. Sagittal sections (50 µm) were then prepared using a vibratome. Sections were mounted on glass slides in 0.1 M PB, cover-slipped, and fluorescence images were acquired using an Olympus SliceView VS200 slide scanner equipped with a Hamamatsu Orca-Fusion camera and a 40× Olympus UPlanXApo objective. Sections including striatal regions enriched in clusters of tdTomato fluorescence were selected for further FIB-SEM processing and processed in sequence through the following steps: incubation in 2.5% glutaraldehyde, 2 mM $CaCl_2$ (J.T. Baker) in 0.1 M sodium cacodylate buffer for 2–3 hours on ice; incubation in 2% osmium tetroxide, 1.5% potassium ferrocyanide ($K_4Fe(CN)_6$) (Sigma-Aldrich), and 2 mM $CaCl_2$ in

0.1 M sodium cacodylate buffer for 1 hour on ice; incubation with thiocarbohydrazide (TCH) for 20 minutes at room temperature; a second incubation in 2% osmium tetroxide in water for 30 minutes at room temperature; incubation in 2% aqueous uranyl acetate overnight at 4 °C; dehydration through a graded ethanol series (20%, 50%, 70%, and 90%; 5 minutes each), followed by three changes of 100% ethanol for 5 minutes each. Sections were then transferred to propylene oxide for 10 minutes at room temperature and infiltrated with Epon resin (Embed 812) in propylene oxide at 25% (several hours), 50% (several hours), and 75% (overnight), followed by 100% Epon overnight and fresh 100% resin for several additional hours. Finally, sections were mounted onto 12-mm glass coverslips in a drop of Epon, excess resin was removed, and samples were transferred to 60 °C oven for 48 hours to induce epon polymerization.

Coverslips with Epon-embedded brain sections were glued onto scanning electron microscope (SEM) aluminum sample-mounting stubs, and platinum *en bloc* coating on the sample surface was carried out with a sputter coater (Ted Pella, Inc.). SEM imaging of the surface of the sections was performed to define regions to be analyzed with FIB-SEM by aligning major tissue landmarks, such as blood vessels, cell bodies and axon bundles with the same structures as observed by previous fluorescence microscopy of the same sections in areas enriched in dystrophic DAergic axon (large clusters of tdTomato fluorescence. FIB-SEM imaging of these regions was performed using a Crossbeam 550 FIB-SEM workstation operating with SmartSEM (Carl Zeiss Microscopy GmbH) and the Atlas 5 engine (Fibics Incorporated). The imaging resolution was set at 8 nm/pixel in the x, y axis with milling being performed at 4 nm/step along the z axis (binned down by 2 when images were exported) to achieve an isotropic resolution of 8 nm/voxel. Images were aligned and exported with Atlas 5 (Fibics Incorporated), further processed, and 3D reconstructed with DragonFly Pro software (Comet Technologies Canada Inc). Except when noted, all reagents were from Electron Microscopy Sciences. A detailed protocol is found at: https://dx.doi.org/10.17504/protocols.io.5jyl84x49g2w/v1.

**FIBSEM at HHMI Janelia Research Campus**. Eight month old female mice C57BL/6;129 were anesthetized with a ketamine/xylazine anesthetic mixture before perfusion with 2% depolymerized paraformaldehyde and 2.5% glutaraldehyde in 0.1 M sodium cacodylate buffer and postfixed in 2% OsO4, 1.5% K4Fe(CN)6, 0.1 M sodium cacodylate buffer 1 h. Subsequently, specimens were stained in block with 2% aqueous uranyl acetate (1 h), dehydrated in increasing concentrations of ethanol, and embedded in Epon. Epon-embedded samples (less than 1 mm in all dimensions) were mounted on sample studs and their tops were trimmed to a 40–100- × 100–300-nm dimension. Samples were then re-embedded in durcupan (EMS, Catalog #14040) by dabbing gently on the trimmed top of the Epon-embedded sample with a small volume of uncured durcupan and then exposing them to 60 °C. Next, cured samples were coated with a thin layer of 10-nm gold and 100-nm carbon. For FIB-SEM imaging, a custom built FIB-SEM, with an FEI Magnum FIB column mounted onto a Zeiss Merlin SEM, was used. SEM imaging was performed using a 200-pA electron beam of 700 V landing energy at 200 kHz with 2-nm spacing in the *x* and *y* directions. A 7-nA 30-kV gallium ion beam was used to remove 2-nm of material in the *z* axis after each SEM image to generate an 8-nm isotropic voxel image stack by an InLens detector (Zeiss electron microscope) in which secondary electrons were filtered out through sample biasing so that only backscattered electrons were detected. For the generation of the 3D reconstructions, membrane contours were traced semi-manually by using 3dmod software. A detailed procedure is available at: https://doi.org/10.25378/janelia.24222688 and in Pang et al. 2023 (2). See also (3).

**Conventional TEM imaging**. Striatal sections prepared as described above for FIBSEM (YALE CCMI facility) were cut with a Leica ultramicrotome to generate 60 nm ultrathin sections. Sections were post-stained with uranyl acetate substitute (Uranyless, EMS), followed by a lead citrate (EMS) solution. Sections were observed in a Talos L 120 C TEM microscope at 80 kV and images were taken with Velox software and a 4k × 4 K Ceta CMOS Camera (Thermo Fisher Scientific). A detailed procedure is available at: https://dx.doi.org/10.17504/protocols.io.36wgqxx65lk5/v1.

**Ex vivo slice preparation for the analysis of DA release.** Mice were terminally anesthetized with a mixture of ketamine (50 mg/kg) and xylazine (4.5 mg/kg) and transcardially perfused with ice-cold modified artificial cerebrospinal fluid (aCSF) ("slicing solution," containing 49.14 mM NaCl, 2.5 mM KCl, 1.43 mM NaH2PO4, 25 mM NaHCO3, 25 mM glucose, 99.32 mM sucrose, 10 mM MgCl2, and 0.5 mM CaCl2); after decapitation, the brain was removed and sectioned in 220-µm-thick coronal slices using a vibratome (VT1200S, Leica Microsystems) in the same slicing solution. Slices were transferred into a holding chamber containing recording aCSF (containing 135.7 mM NaCl, 2.5 mM KCl, 1.25 mM NaH2PO4, 25 mM NaHCO3, 2 mM CaCl2, 1 mM MgCl2, and 3.5 mM glucose). Slices were allowed to recover for 30 min at 34°C and then kept at room temperature for at least 15 min before starting the experiments. All solutions were pH 7.4 and ~310 mOsm and were continually bubbled with 95% O2 and 5% CO2. A detailed procedure is available at: https://dx.doi.org/10.17504/protocols.io.ewov1rb8ylr2/v2.

**Two-photon laser scanning microscope (2PLSM) workstation.** The 2PLSM workstation used was an Ultima dual-excitation-channel scan head (Bruker Nano Fluorescence Microscopy Unit). The foundation of the system is the Olympus BX-51WIF upright microscope with an LUMPFL 60×/1.0 NA (numerical aperture) water-dipping objective lens. The automation of the XY stage motion, lens focus, and manipulator XYZ movement was provided by an FM-380 shifting stage, an axial focus module for Olympus scopes, and manipulators (Luigs & Neumann). The 2P excitation (2PE) source was a Chameleon Ultra1 series tunable wavelength (690 to 1040 mm, 80 MHz, ~250 fs at sample) Ti: sapphire laser system (Coherent Laser Group). Each imaging laser output is shared (equal power to both sides) between two optical workstations on a single anti-vibration table (TMC). Workstation laser power attenuation was achieved with two Pockels' cell electro-optic modulators (models M350-80-02-BK and M350-50-02-BK, Con Optics) controlled by Prairie View 5.6 software. The two modulators were aligned in series to provide enhanced modulation range for fine control of the excitation dose (0.1% steps over 5 decades), to limit the sample maximum power. The fluorescence emission was collected by non–de-scanned photomultiplier tubes (PMTs). Green channel (490 to 560 nm) signals were detected by a Hamamatsu H7422P-40 select GaAsP PMT. Red channel (580 to 630 nm)

signals were detected by a Hamamatsu R3982 side on PMT. Dodt-tube–based transmission detector with Hamamatsu R3982 side on PMT (Bruker Nano Fluorescence) allowed visualization of slices during laser scanning. Scanning signals were sent and received by the National Instruments PCI-6110 analog-to-digital converter card in the system computer (Bruker Nano Fluorescence). All spiral images were collected with a pixel size of 0.393 µm, a pixel dwell time of 8 µs, and a frame rate of 39.165 fps (frames per second).
A detailed protocol is found at:
https:// dx.doi.org/10.17504/protocols.io.bp2l6jwn1vqe/v1.

**DA release measurements.** Mice homozygous or heterozygous for the RQ mutation, referred to as $Synj1^{RQ}$ mice and *Synj1*$^{+/RQ}$ or control mice, respectively, were stereotaxically injected with 500 nl of AAV5-CAG-dLight1.3b Addgene #125560) into either the DLS (AP: 0.3, ML: 1.3, DV: 3.1) or NAc (AP: 1.1, ML: 1.1, DV 4.0) at P30. The Allen Mouse Brain Atlas, online version 1, 2008 (http://mouse.brain-map.org) was the source of all stereotaxic coordinates. Experiments were performed at P45 (+/- 2 days). Mice were anesthetized, perfused and brain slices prepared as described above and previously (4). All dLight recordings were performed in the continuous presence of mecamylamine (10µM), sulpiride (10µM) and gabazine (10µM) in the recording aCSF to block nicotinic acetylcholine receptors (5, 6), D2/D3 DA autoreceptors (7), and GABA-A receptors respectively, and thereby isolate electrically evoked DA transients from confounding neuromodulatory influences. dLight imaging was performed with 920 nm light using the 2PLSM system, as defined above. dLight samples were imaged with 20 µm diameter field of view (continuous spiral scan with 95% duty cycle using 84,267 px, 8.0 µs pixel dwell time at 39.165 f.p.s. All ROI signals were summed per frame. The spiral scan data (outside of the circle towards the inside of the circle) has 3000 ms of pre-stimulation baseline ($f_0$, with laser blocked background counts subtracted). DA release was evoked by electrical stimulation (600 µA, 0.2 ms pulse width, 1P or 5 pulses at 10 Hz) using a tungsten bipolar electrode inserted approximately 30 µm to either the DLS orVS In some experiments a stimulation was applied every 10 seconds for 5 times (8, 9). Boxplots quantifying DA release are presented as a percentage of ∆F max (F-max was attained by applying 100 µM DA to the slice), calculated by F-max divided by F0. The time-series plots display $\Delta f/f_0$ normalized to the first response in the series. Traces representing DA release are normalized to ΔF max to allow comparisons between slices and animals. A detailed protocol is found at: https://dx.doi.org/10.17504/protocols.io.bp2l6jwn1vqe/v2 .

**Analysis and statistics**. Images were analyzed offline using FIJI (10). Traces were subsequently analyzed using Excel (Microsoft) and analyzed in GraphPad Prism8 (GraphPad software). Sample n represents the number of brain slices collected from N animals. Statistical analysis was performed using GraphPad Prism8. The non-parametric Mann-Whitney U test was used to assess statistical significance in experiments measuring absolute DA release. For experiments investigating vesicle recycling, a two-way repeated measures ANOVA was used in conjunction with the Šídák method to correct for multiple comparisons. Probability threshold for statistical significance was $P < 0.05$.

**References for Extended Methods**


1. M. Cao *et al.*, Parkinson Sac Domain Mutation in Synaptojanin 1 Impairs Clathrin Uncoating at Synapses and Triggers Dystrophic Changes in Dopaminergic Axons. *Neuron* **93**, 882-896.e885 (2017).
2. S. Pang, C. S. Xu, Methods of enhanced FIB-SEM sample preparation and image acquisition. *Methods Cell Biol* **177**, 269-300 (2023).
3. C. S. Xu *et al.*, Enhanced FIB-SEM systems for large-volume 3D imaging. *Elife* **6** (2017).
4. E. Zampese *et al.*, Ca(2+) channels couple spiking to mitochondrial metabolism in substantia nigra dopaminergic neurons. *Sci Adv* **8**, eabp8701 (2022).
5. H. Zhang, D. Sulzer, Frequency-dependent modulation of dopamine release by nicotine. *Nat Neurosci* **7**, 581-582 (2004).
6. S. Chatterjee *et al.*, The α5 subunit regulates the expression and function of α4*-containing neuronal nicotinic acetylcholine receptors in the ventral-tegmental area. *PLoS One* **8**, e68300 (2013).
7. S. Boumhaouad *et al.*, Regulation of Dopamine Release by Tonic Activity Patterns in the Striatal Brain Slice. *ACS Chem Neurosci* **16**, 303-310 (2025).
8. N. J. Platt, S. Gispert, G. Auburger, S. J. Cragg, Striatal dopamine transmission is subtly modified in human A53Tα-synuclein overexpressing mice. *PLoS One* **7**, e36397 (2012).
9. T. Virmani, D. Atasoy, E. T. Kavalali, Synaptic Vesicle Recycling Adapts to Chronic Changes in Activity. *The Journal of Neuroscience* **26**, 2197-2206 (2006).
10. J. Schindelin *et al.*, Fiji: an open-source platform for biological-image analysis. *Nat Methods* **9**, 676-682 (2012).

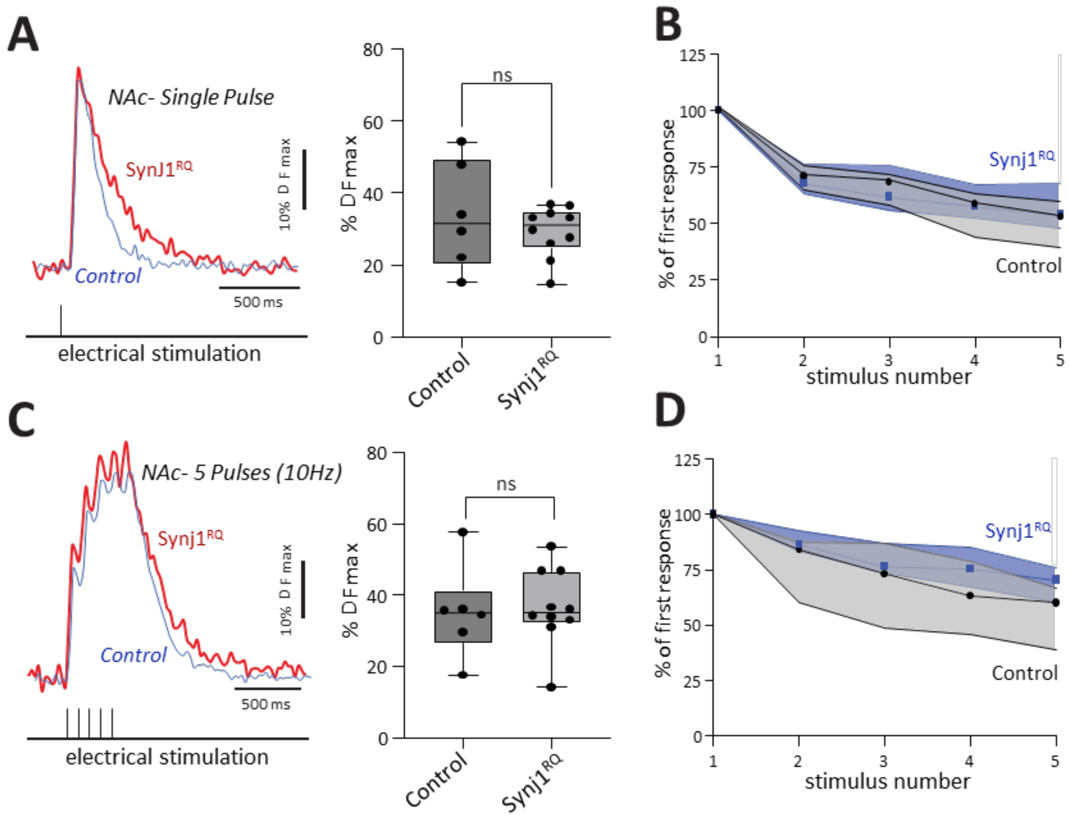


**Fig. S1. No significant defect in evoked DA release in the Nucleus Accumbens (NAc) of the ventral striatum (VS) of Synj1$^{RQ}$ mice. A.** Representative traces of DA release in the NAc evoked by single-pulse stimulation. Traces are normalized to dF max (left). Quantification of the amount of DA released from terminals innervating the NAc in response to single stimulatory events (right). Quantifications are presented as a percentage of dF max (achieved through bath application of 100 µM DA). Synj1$^{RQ}$: n=6, N=3; *Synj1*$^{+/RQ}$ (control): n=10, N=2. **B.** Plot illustrating no difference in evoked DA release when release is evoked by consecutive single pulses. Responses are normalized to the first response in the stimulation protocol. *Synj1*$^{+/RQ}$ (control): n=10, N=2. Synj1$^{RQ}$: n=6, N=3. **C.** Representative traces of DA release in the NAc in response to 5-pulse stimulation at 10 Hz. Traces are normalized to dF max (left). Quantification of the amount of DA released from terminals innervating the NAc in response to a 5-pulse train at 10 Hz. Quantifications are based on the dF of the 5$^{th}$ pulse in the train and are presented as a percentage of dF max (achieved through bath application of 100 µM DA). Synj1$^{RQ}$: n=6, N=2; *Synj1*$^{+/RQ}$ (control): n=10, N=3. (right). **D.** Deficits in vesicle recycling are absent in the NAc of Synj1$^{RQ}$ mice when release is evoked a 5-pulse train at 10 Hz. Responses are normalized to the first response in the stimulation protocol. *Synj1*$^{+/RQ}$: n=10, N=3. Synj1$^{RQ}$: n=6, N=2.

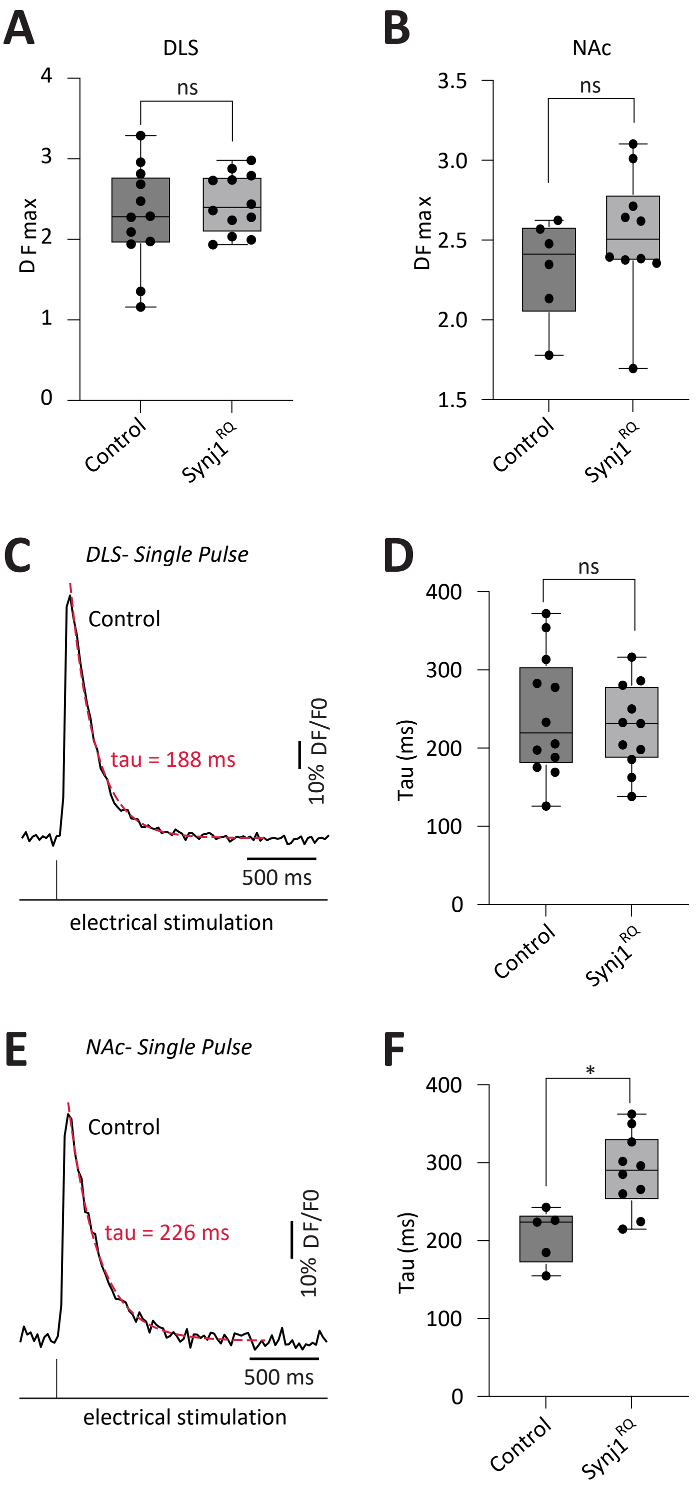
A
DLS
ns
D F max
Control
Synj1RQ
B
NAc
ns
D F max
Control
Synj1RQ
C
DLS- Single Pulse
Control
tau = 188 ms
10% DF/F0
500 ms
electrical stimulation
D
ns
Tau (ms)
Control
Synj1RQ
E
NAc- Single Pulse
Control
tau = 226 ms
10% DF/F0
500 ms
electrical stimulation
F
*
Tau (ms)
Control
Synj1RQ

**Fig. S2. Analysis of dLight signal DF max and decay in different genotypes. A.** Boxplot of the maximum fluorescence response (ΔF max) evoked by bath application of 100µM DA in the DLS of *Synj1*$^{+/RQ}$ (control) and Synj1$^{RQ}$ mice, used as the normalization reference for all dLight1.3b recordings in this region. No significant difference in ΔF max was detected between genotyes, confirming comparable dLight1.3b expression levels and sensor dynamic range across groups. *Synj1*$^{+/RQ}$ (control): n=12, N=7; Synj1$^{RQ}$: n=12, N=6. **B**. Boxplot of the maximum fluorescence response (ΔF max) evoked by bath application of 100µM DA in the NAc of *Synj1*$^{+/RQ}$ (control) and Synj1$^{RQ}$ mice, used as the normalization reference for all dLight recordings. No significant difference in ΔF max was detected between genotypes, confirming that dLight1.3b expression levels and sensor dynamic range are comparable across groups. *Synj1*$^{+/RQ}$ (control): n=10, N=2; Synj1$^{RQ}$: n=6, N=3. **C.** Representative single-pulse evoked dLight1.3b fluorescence trace (black) from a *Synj1*$^{+/RQ}$ mouse (control) with a single-exponential decay function fitted to the post-peak signal (dashed red line; $\tau$ = 188ms). Scale bars: 10% ΔF/F0 (vertical), 500 ms (horizontal). **D.** Boxplot of decay time constants ($\tau$, ms) for all recorded DLS transients across genotypes. No significant difference was detected between *Synj1*$^{+/RQ}$ (control) and Synj1$^{RQ}$ animals (Mann-Whitney U, p = 0.7859, n = 12 and n = 11, respectively). **E.** Representative single-pulse evoked dLight1.3b fluorescence trace (black) from a *Synj1*$^{+/RQ}$ mouse (control) with a single-exponential decay fit overlaid (dashed red line; $\tau$ = 226 ms). Scale bars: 10% ΔF/F0 (vertical), 500 ms (horizontal). **F.** Boxplot of decay time constanys ($\tau$, ms) across genotypes in the NAc. Synj1$^{RQ}$ animals exhibited significantly prolonged DA clearance kinetics compared to *Synj1*$^{+/RQ}$ control littermates (Mann-Whitney U, P = 0.0127, n = 10 and n = 5 slices, respectively).

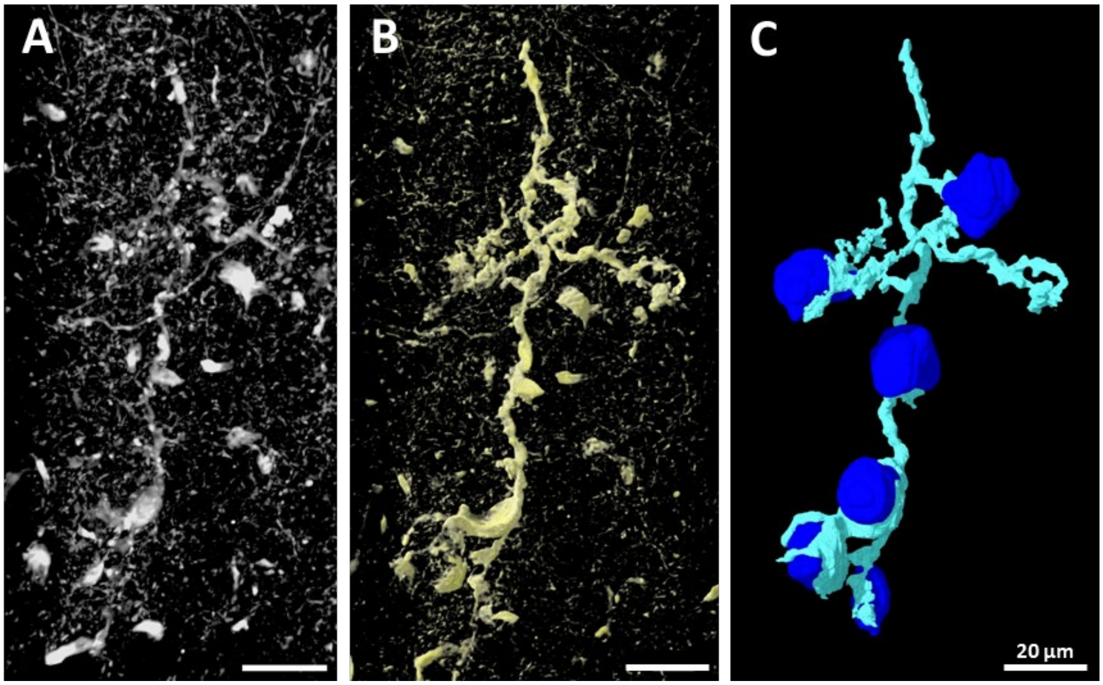


**Fig. S3**. **cell bodies in the DLS of a delipidated tdTomato-Synj1$^{RQ}$ mouse brain. A.** Confocal imaging of a 100 µm thick vibratome section of delipidated DLS from a Synj1$^{RQ}$ mouse expressing tdTomato in DAergic axons. The large dystrophic DA axon stands out among the fine network of normal DA axons. **B** and **C**. 3 dimensional views of the large dystrophic axon shown in A. **C**. Segmentation of the axon (cyan) shown in A and B, revealing its contacts with 6 cell bodies (dark blue). Scale bars: 20 µm.

**Legends for movies**

**Movie S1** (separate file). **3D view of dystrophic DAergic axons distribution in the striatum of a Synj1$^{RQ}$ mouse expressing tdTomato in DAergic neurons**. The entire mouse brain had been cleared by delipidation and the tdTomato fluorescence (white) was imaged by light sheet microscopy. Only the striatum, defined by the fluorescence of tdTomato, is shown. Large foci of tdTomato fluorescence were segmented and shown in red superimposed onto the diffuse (white) tdTomato fluorescence corresponding to the normal network of DAergc axons. Note the selective concentration of the large fluorescence foci (dystrophic axon segments) in the dorsal striatum. A small volume of the region most enriched in dystrophic axons is shown at higher magnification in Movie S3. The scattered white spots visible in the VS represent cell bodies expressing low levels of tdTomato probably as an off-target effect of the DAT promoter used for the study. They stand out in this high contrast movie.

**Movie S2** (separate file). Sequence (rostral to caudal) of coronal views of optical section of the striatum shown in Fig. 1D and in movie S1.

**Movie S3** (separate file). **3D view of a small volume of the region most enriched in dystrophic axons from the striatum shown in movie S1 and S2.** A 100 µm thick vibratome section was generated from the cleared brain and examined by confocal microscopy. A snapshot of this movie is shown in Fig. 1F. Scale bar: 30 µm.

**Movie S4** (separate file). **FIB-SEM stack of a small volume of the striatum of a Synj1$^{RQ}$ mouse obtained with a Zeiss Crossbeam 550 (Yale CCMI).** The movie illustrates the two DAergic dystrophic axon segments numbered #2 and #3 in Fig. 4. An astrocyte is pseudocolored in light green. Dystrophic axons are pseudocolored in light magenta and mitochondria in the dystrophic axons in dark green. (Scale bar: 1.5µm)

**Movie S5** (separate file). **3D reconstruction of the FIB-SEM volume of the DAergic dystrophic axon segments numbered #2 and #3 in Fig. 4.** Green = surface of an astrocyte; blue = blood vessel lumen; pink = surface view of DAergic dystrophic axon segments; magenta = membrane whorls, dark green = mitochondria. Figure 3I-L are snapshots of this movie. (Scale bar: 1 µm)

**Movie S6** (separate file). **FIB-SEM stack of a small volume of the striatum of a Synj1$^{RQ}$ mouse at 8 x 8 x 8 nm voxel size with a customized equipment (HHMI Janelia Research Campus).** The movie shows that onion-like structures surround the evaginations of a neighboring cell. A red line outlines the outer surface of the dystrophic axon. A cyan line outlines the plasma membrane of the evaginations of the neighboring cell trapped into the dystrophic axon. (Scale bar: 800nm)

**Movie S7** (separate file). **3D reconstruction of the structures shown in Movie S6.** Only the outer surface of the dystrophic axon terminal (light blue), the membrane whorl (brown) and the invagination of the neighboring cell are shown. (Scale bar: 800nm)

**Movie S8** (separate file). **Z stack images of the field shown in Fig. S2A and B.** The tdTomato fluorescence is shown in white and the large dystrophic axon that contacts the cell bodies (dark blue) is pseudocolored in cyan.

**Movie S9** (separate file). **3D view of the image stack shown in Movie S8**.

**Movie S10** (separate file). **3D reconstruction of the large dystrophic DAergic axon shown in Fig. S2B and C and in movies S8 and S9.** Yellow = saturated fluorescence intensity of tdTomato in the dystrophic DAergic axon; cyan = segmented dystrophic axon; blue = cell bodies contacted by the dystrophic DAergic axon.
